# Recognition of unique cell surface glycopolymers by diversified lectin domains specifies umbrella toxin targeting

**DOI:** 10.1101/2025.04.24.650519

**Authors:** Qinqin Zhao, Jiri Vlach, Young-Jun Park, Yongjun Tan, Savannah K. Bertolli, Pooja Srinivas, Pinyu Liao, Connor R. Fitzpatrick, Jeffery L. Dangl, Parastoo Azadi, Frank DiMaio, S. Brook Peterson, Dapeng Zhang, David Veesler, Joseph D. Mougous

**Author notes:** To whom correspondence should be addressed: Email –.

## Abstract

The bacterial cell envelope provides structural integrity and is an essential conduit through which the organism interacts with its environment. However, the molecular structures of its constituents can also be exploited as receptors, permitting threats such as toxins and phage access to the cell. We previously demonstrated that *Streptomyces coelicolor* secretes an umbrella toxin particle that acts in a highly selective manner to inhibit the hyphal growth of competing *Streptomyces* strains. Here, we identify the receptor of umbrella toxins as teichuronic acid (TUA) oligosaccharides anchored to the cell surface through linkage to wall teichoic acids (WTA). We show that the carbohydrate portion of this previously undescribed hybrid TUA–WTA molecule is variable across species and that targeting specificity derives from its selective recognition by diversified lectin domains associated with umbrella particles. A cryo-EM structure of a lectin–TUA complex reveals the molecular basis for umbrella toxin cell targeting and, in conjunction with bioinformatic analyses, provides insights into a molecular arms race that we posit drives diversification of umbrella particles and the chemical composition of *Streptomyces* cell envelopes.

## Introduction

The bacterial cell surface is an evolutionary battleground. Its constituents serve vital roles for the cell, yet these molecules are also vulnerable to exploitation as receptors for phage, immune system effectors, and antibacterial toxins. Extracellular carbohydrates, lipopolysaccharide, teichoic acids, protein appendages such as pili and flagella, and outer membrane proteins are all subject to such exploitation^1–4^. The evolutionary trajectory of these molecules is thus driven by opposing pressures: selection for diversification to escape receptor recognition, and stabilizing selection to maintain functionality.

*Streptomyces* are a genus of bacteria known for producing antimicrobial small molecules^5–7^. As ubiquitous inhabitants of soil and other habitats rich in microbial diversity, they deploy this arsenal of antimicrobials to deter a wide array of microorganisms^6^. This is particularly critical when *Streptomyces* transition from vegetative growth to spore formation; nutrients released during programmed cell death at this stage support spore development and require protection from opportunistic microbes^8^. *Streptomyces* also experience significant competition from other members of their genus. Experiments routinely show the co-existence of multiple *Streptomyces* species in soil, highlighting the ecological relevance of this congeneric competition^9^. However, the consequences of this form of competition on the evolution of *Streptomyces* physiology are largely unknown.

We recently reported the discovery that *Streptomyces* elaborate complex, proteinaceous toxigenic particles that mediate congeneric competition^10^. We named these umbrella toxins based on their morphology, and demonstrated that one of the three such toxins from *S. coelicolor*, the Umb2 particle, acts in a highly selective manner to inhibit hyphal growth of other Streptomyces species. Umbrella toxin particles are composed of at least three distinct polypeptides, UmbA-C.

UmbC is a large protein that forms the core of the particle and harbors the domain conferring toxicity. The protein consists of a ring composed of eight degenerate repeats on one end, connected to the toxin domain on the other end by an extended coiled-coil stalk. Five repeats of the UmbC ring are bound by UmbB, a small adaptor protein able to concurrently bind UmbA. UmbA–B complexes thereby form five spokes emanating from the UmbC ring, lending the particle its umbrella moniker. UmbA contains two domains: an N-terminal trypsin-like domain that mediates UmbB binding and a C-terminal lectin-like domain of unknown function. *S. coelicolor* possesses six UmbA proteins (UmbA1-6); UmbA1-3 associate with single Umb particles, Umb1-3, respectively, whereas the other three bind promiscuously to all three (Figure 1A).

**Figure 1.**
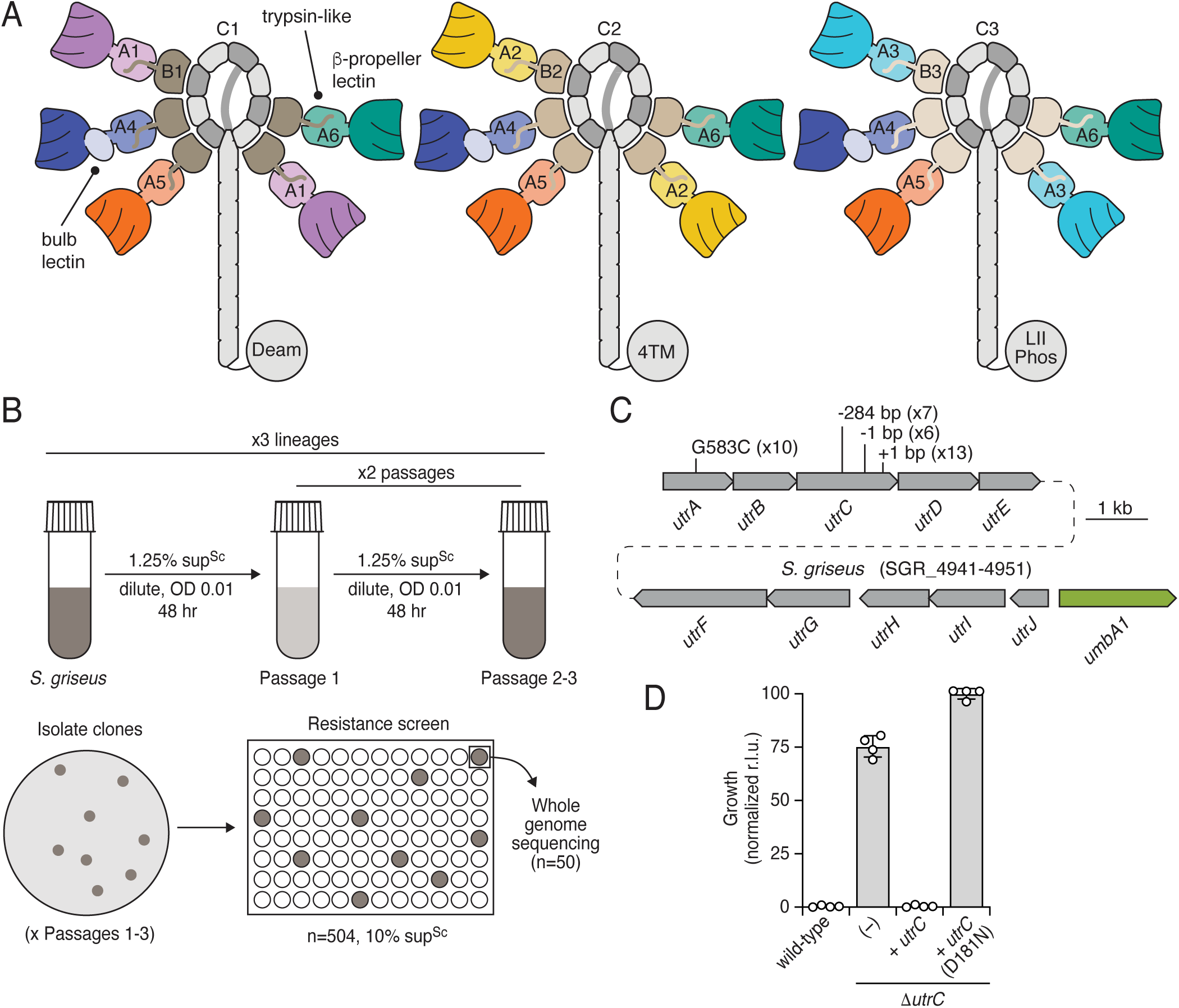
Mutations in a candidate carbohydrate biosynthesis operon of *S. griseus* confer umbrella toxin resistance. A) Schematic illustrating the composition of the three umbrella toxin particles produced by *S. coelicolor.* The complement of UmbA proteins associated with a given particle can vary; schematics depict those specialized for a given particle (UmbA1-3) and those which can be present on each (UmbA4-6). Toxin domains are labeled according to their predicted activity or structure: Deam, deaminase; 4TM, four transmembrane helices; LII Phos, lipid II phosphatase. Colors are consistent with depictions of these molecules in other figures. B) Overview of adaptive evolution (ALE) experiment to isolate *S. griseus* clones resistant to *S. coelicolor* umbrella toxins. C) Schematic depicting the location of mutations in an *S. griseus* carbohydrate biosynthesis gene cluster selected during ALE with sup^Sc^ treatment. Numbers indicate the number of times the indicated mutation was observed across 50 clones sequenced. The location of the sole *umbA* gene in *S. griseus* is also indicated. D) Growth yields of the indicated strains of *S. griseus* after 16 h of treatment with sup^Sc^.

Prior bioinformatic analyses suggest that umbrella particles are produced by a significant proportion of *Streptomyces* species, as well as many other bacteria belonging to the Actinomycetota phylum^10^. These findings also highlighted sites of hypervariability within umbrella particles. One such site is the polymorphic toxin domain of UmbC proteins, a common characteristic of interbacterial antagonism machineries that reflects the evolution of resistance among other selective pressures. A surprising site of hypervariability in umbrella particles occurs within the lectin-like domains of UmbA proteins. Cumulatively, UmbA proteins possess at least 14 structurally distinct predicted lectins, often with a single protein carrying more than one.

Moreover, there is extensive sequence variation within these large fold families, suggestive of differential binding specificity. This variability is compounded by the observation that multiple UmbA proteins can associate with a single umbrella toxin particle. The function of UmbA proteins was not determined, but it was speculated that these proteins – via their lectin-like domains – might engage receptors on target cells.

Regardless of the cell type they target, protein toxins secreted by bacteria require specific interactions with receptor molecules on target cell surfaces^11^. This specificity avoids off-target adsorption, offsetting the high cost of their biosynthesis and export. For example, colicins achieve strict intraspecies targeting by binding to conserved proteins embedded in the outer-membrane^4^. These specific interactions also facilitate the evolution of resistance, leading to molecular arms races between producers and targets that can have far reaching consequences ^12^. Here, we show that UmbA proteins mediate receptor binding on target *Streptomyces* species. We identify the receptor as a previously undescribed glycoconjugate consisting of a teichuronic acid (TUA) polymer appended to wall teichoic acids (WTA) and use cryo-EM to elucidate the structural basis for its recognition by a UmbA lectin. We show that variation in TUA structure mediates differential targeting by umbrella toxins, and provide bioinformatic evidence of a molecular arms race driving diversification of the umbrella toxin lectins and the carbohydrates decorating the *Streptomyces* cell surface that they exploit.

## Results

### Identification of a carbohydrate biosynthetic gene cluster implicated in umbrella toxin resistance

We previously showed that the Umb particles of *S. coelicolor* act in a highly selective manner to intoxicate other *Streptomyces* spp, including *S. griseus*^10^. We hypothesized that this selectivity could derive, at least in part, from the variable presence of cognate receptor molecules on target cells. However, the molecular identity of a Umb particle receptor has not been determined. As a first step toward defining a Umb particle receptor, we selected for *S. griseus* strains resistant to intoxication by high molecular weight protein-enriched *S. coelicolor* supernatant (sup^Sc^) (Figure 1B). Briefly, we propagated three independent lineages of *S. griseus* at a concentration of sup^Sc^ sufficient to inhibit ∼90% of growth after 16 hrs (Figure S1A). After three passages, we observed robust *S. griseus* growth in the toxin-treated samples, approximating that of the untreated control. Analysis of individual clones isolated from each passage revealed that only 6% (28/432) of them exhibited sup^Sc^ resistance after one passage, which increased to 35% (17/48) and 100% (24/24) after two and three passages, respectively (Figure S1B).

Whole genome sequencing of 50 resistant clones deriving from all passages and lineages revealed 11 genes with mutations in multiple clones (Table S1). Among these, those within a single previously uncharacterized gene cluster encompassing open reading frames (ORFs) SGR_4941-4945 were most prevalent. We named these ORFs umbrella toxin resistance genes A-E (*utrA*-*E*) (Figure 1C). Non-synonymous and assorted frameshift mutations in *utrA* or *utrC* were observed in each lineage and 72% of all sequenced clones. In many cases, a mutation in this region accounted for the only non-synonymous mutation in a given clone, strongly suggesting a causative role in sup^Sc^ resistance (Table S1). To evaluate this directly, we generated a *S. griseus* strain bearing an in-frame deletion of *utrC*. Like strains selected by toxin exposure, the growth of this strain was unaffected by sup^Sc^ at a level sufficient to fully inhibit growth of the wild-type (Figure 1D). Plasmid-based complementation of the intact gene reverted the strain to the Umb-sensitive phenotype.

Bioinformatic analyses of UtrC revealed that the protein possesses a predicted glycosyltransferase domain distantly resembling that found in *S. aureus* TarS, a protein that installs β-linked N-acetylglucosamine on the ribitol phosphate backbone of wall teichoic acids (WTAs) (Figure S1C) ^13^. This domain includes a predicted catalytic aspartate and we found that substitution of the corresponding residue in UtrC with asparagine (D181N) abolishes sup^Sc^ sensitivity, presumably via abrogating its function^14^ (Figure 1D). Interestingly, further analyses of the *utrA-E* cluster indicated that all five of the genes encode proteins strongly predicted to act on carbohydrates (Table S2). For example, *utrA*, the other gene hit in our selection, encodes a likely sugar aminotransferase and *utrE* encodes a predicted epimerase. Also of note, immediately downstream of this gene cluster is a divergently transcribed operon of five genes, which we renamed *utrF-J*, encoding additional predicted carbohydrate modifying enzymes (Table S2). One of these, UtrF shows strong predicted structural similarity to *S. aureus* TarM, which installs α-linked N-acetylglucosamine on the ribitol phosphate backbone of WTAs^15^. Adjacent to and divergently transcribed from *utrJ* is the ORF encoding the sole *S. griseus* UmbA ortholog, further suggesting a link between the *utr* genes and umbrella toxin-based antagonism.

### Synthesis of a TUA–WTA hybrid polymer is required for Umb-based intoxication

Given the bioinformatic links between multiple Utr proteins and WTA biosynthetic enzymes, we sought to compare WTA structures of *S. griseus* wild-type and Δ*utrC* strains. WTAs from these strains were obtained by hydrolysis from isolated cell wall sacculi as described previously, with minor modifications^16^. Analysis of NMR spectra of the wild-type preparation revealed ^1^H-^13^C HSQC peaks corresponding to carbohydrate and polyol phosphate (polyol*P*) moieties, consistent with carbohydrate-modified WTA structures present in related organisms (Figure 2A)^17^. Strikingly, while the Δ*utrC* preparation contained similar polyol*P* peaks, those attributable to carbohydrates were either strongly diminished or undetected. This finding suggests that the glycosyltransferase encoded by *utrC* is required for the biogenesis of the predominant carbohydrate modification to the WTA of *S. griseus*.

**Figure 2.**
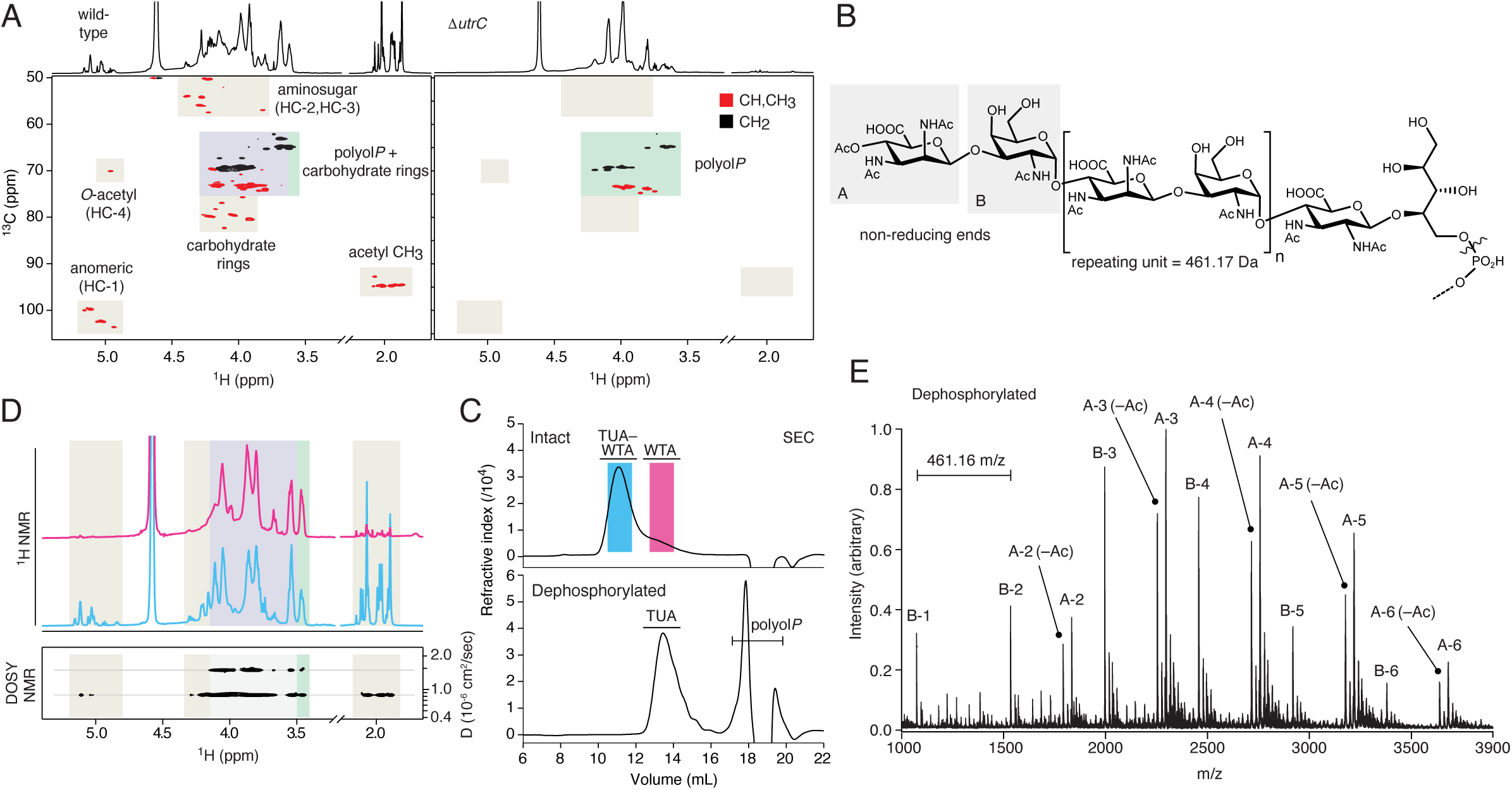
Structure of a TUA–WTA hybrid polymer produced by *S. griseus* and required for targeting by *S. coelicolor* umbrella toxins. A) Partial views of 2D multiplicity-edited ^1^H-^13^C HSQC NMR spectra of *S. griseus* wild-type (left) and Δ*utrC* (right) intact WTA polymers obtained by hydrolysis from isolated cell wall sacculi. The corresponding 1D ^1^H spectra are shown above each HSQC panel. Grey shading indicates peaks deriving from carbohydrates; green shading indicates poloyl*P*-derived peaks; blue shading indicates peaks deriving from both polyol*P* moieties and carbohydrates. B) Structure determined for the *S. griseus* TUA →[4)-ManNAc3NAcA-β(1→3)-GalNAc-α(1→]_n_ oligomer. ∼70% of the TUA is terminated with a 4OAc-ManNAc3NAcA residue (non-reducing end A), while ∼30% is terminated with a GalNAc residue (non-reducing end B). The reducing-end residue of GlcNAc3NAcA forms a β(1→4) glycosidic bond with a ribitol-5-phosphate residue. The dashed bond indicates the expected linkage with the polyol-*P* WTA polymer; the wavy line indicates the site of cleavage upon chemical dephosphorylation. C) SEC analysis of intact (top) or dephosphorylated polymer obtained from *S. griseus* WT cell wall preparations by TCA hydrolysis. D) ^1^H NMR (top) and DOSY (bottom) spectra of material from the two SEC peaks obtained for the total cell wall polymer isolate. NMR peak colors correspond to panel (C), shading corresponds to panel (A). D, diffusion coefficient. E) MALDI-TOF mass spectrometry analysis of dephosphorylated polymer obtained from wild-type *S. griseus.* Masses corresponding to [M−H]^−^ ions of TUA oligomers with reducing ends A (2–6 repeating units) and B (1–6 repeating units) were detected. The signals of the acetylated reducing end A oligomers were accompanied by secondary peaks due to the loss of acetyl group.

Detailed analyses of our 2D NMR unexpectedly revealed that the carbohydrate molecule present in wild-type samples and missing from Δ*utrC* consists of an extended glycan polymer with a repeating unit of →3)GalNAc-α(1→4)-ManNAc3NAcA-β(1→, alternately terminated at the non-reducing end by 4-OAc-ManNAc3NAcA or GalNAc residues (Figure 2B, Figure S2A and Table S3). The repeating portion of this structure is reminiscent of a distinct and poorly characterized class of cell wall-associated anionic polymers called teichuronic acids (TUAs), found in *Streptomyces* and several other genera of Actinomycetota and Bacillota^17,18^.

These polysaccharide molecules are tacitly defined by the presence of a uronic acid that forms 2–4-membered repeating units with neutral monosaccharides. In certain species, including *Micrococcus luteus* and *Bacillus licheniformis*, there is evidence for direct linkage of TUAs to the cell wall by a phosphodiester bond to the *N*-acetylmuramic acid sugar of peptidoglycan^19,20^. However, our analyses indicate that in the *S. griseus* TUA-like molecule we identified, the reducing-end sugar, GlcNAc3NAcA, is directly linked by a β glycosidic bond to the 4-hydroxyl of ribitol 5-phosphate (Figures 2B, S2A, and Table S3). Additionally, peaks in the ^1^H-^13^C HSQC spectra not attributable to this TUA-like molecule derive from 1,5- and 1,3,5-linked ribitol*P* units, as well as a terminal 3-linked glycerol*P* moiety, suggestive of a mixed-type ribitol/glycerol WTA^21,22^. In contrast, the equivalent polymer from *S. griseus* Δ*utrC* lacked the 1,3,5-linked ribitol*P*, and contained an additional terminal 5-linked ribitol*P* residue reminiscent of the 5-linked ribitol*P* that is substituted with the TUA-like molecule in wild-type *S. griseus*. These findings raised the intriguing possibility that in *S. griseus*, and perhaps other *Streptomyces* species, TUA polymers are appended to WTAs. Moreover, given the resistance of *S. griseus* Δ*utrC* to umbrella toxins, such a hybrid TUA–WTA molecule could serve as a receptor for umbrella toxin particles on the cell surface.

To begin evaluating these possibilities, we performed size exclusion chromatography on the material released from a wild-type *S. griseus* cell wall preparation. This revealed the presence of two peaks (Figure 2C); analysis of fractions representing each peak by ^1^H and ^1^H-^13^C HSQC NMR provided evidence that the higher mass species constitutes a polymer with both the TUA-like and WTA components, whereas the lower mass peak contains only those indicative of WTAs (Figure 2D, Figure S2B and Table S3). Diffusion-ordered NMR spectroscopy (DOSY) showed that the *S. griseus* TUA and WTA polymers have the same hydrodynamic properties in the higher-mass species, indicative of a single molecule (Figure 2D). The complexity of ^1^H, ^13^C and ^31^P spectra prevented us from establishing a direct link between the TUA and WTA by NMR. However, dephosphorylation of the material yielded a major species with significantly reduced molecular weight, consistent with degradation of the polyol*P* WTA backbone (Figure 2C). Analysis of this material by NMR and mass spectrometry confirmed its structure as that of the intact TUA portion of the glycoconjugate (Figure 2E, Figure S2C and Table S3). Together, these data indicate that UtrC is required for biosynthesis of an extended, TUA-like polymer linked to a *S. griseus* WTA backbone (Figure S2D,E).

Though the TUAs of numerous *Streptomyces* sp, including *S. griseus*, are chemically characterized, the genes involved in their biosynthesis have remained unidentified^17^. Indeed, *B. subtilis* is the only bacterium for which the genetic basis of TUA biosynthesis is known^23,24^. Based on predictions and limited supporting biochemical data, *B. subtilis* is thought to synthesize a TUA polymer disaccharide subunit intracellularly on the undecaprenyl lipid carrier. Following export by TuaB, a predicted membrane flippase, and extracellular polymerization with additional subunits by TuaE and TuaF, the TUA is transferred from the lipid carrier to peptidoglycan^24^. The *S. griseus* TUA operon lacks equivalents of the membrane associated, extracellular polymerase enzymes required for TUA biosynthesis in *B. subtilis* TUA, nor does it encode a membrane flippase. The functions encoded within the TUA operon of this species instead point toward direct, intracellular modification of WTA.

### UmbA4 associates with *S. griseus* in a TUA-dependent manner

The lectin domains of UmbA proteins are positioned at the tips of the five spokes of Umb particles^10^. Our finding that a cell surface carbohydrate is required for Umb-based intoxication led us to speculate that these proteins might function as the receptor binding modules of umbrella toxins. To test this, we sought to measure the capacity of individual *S. coelicolor* UmbA proteins to mediate target cell binding. Prior work by our laboratory demonstrated that the Umb2 particle of *S. coelicolor* is solely responsible for the inhibitory activity of sup^Sc^ against *S. griseus*^10^. Given that *S. coelicolor* Umb particles possess a heterogenous assortment of UmbA proteins (e.g. Umb2 contains UmbA2 and UmbA4-6), assessing whole particle binding was deemed an impractical solution. However, we also found UmbA proteins generally challenging to produce and purify heterologously. To circumvent these potential limitations, we used sup^Sc^ from a strain lacking UmbC1-3 as a source of the UmbA proteins. In this background, the UmbA proteins are untethered from each other, allowing their individual binding specificity to be ascertained (Figure 3A).

**Figure 3.**
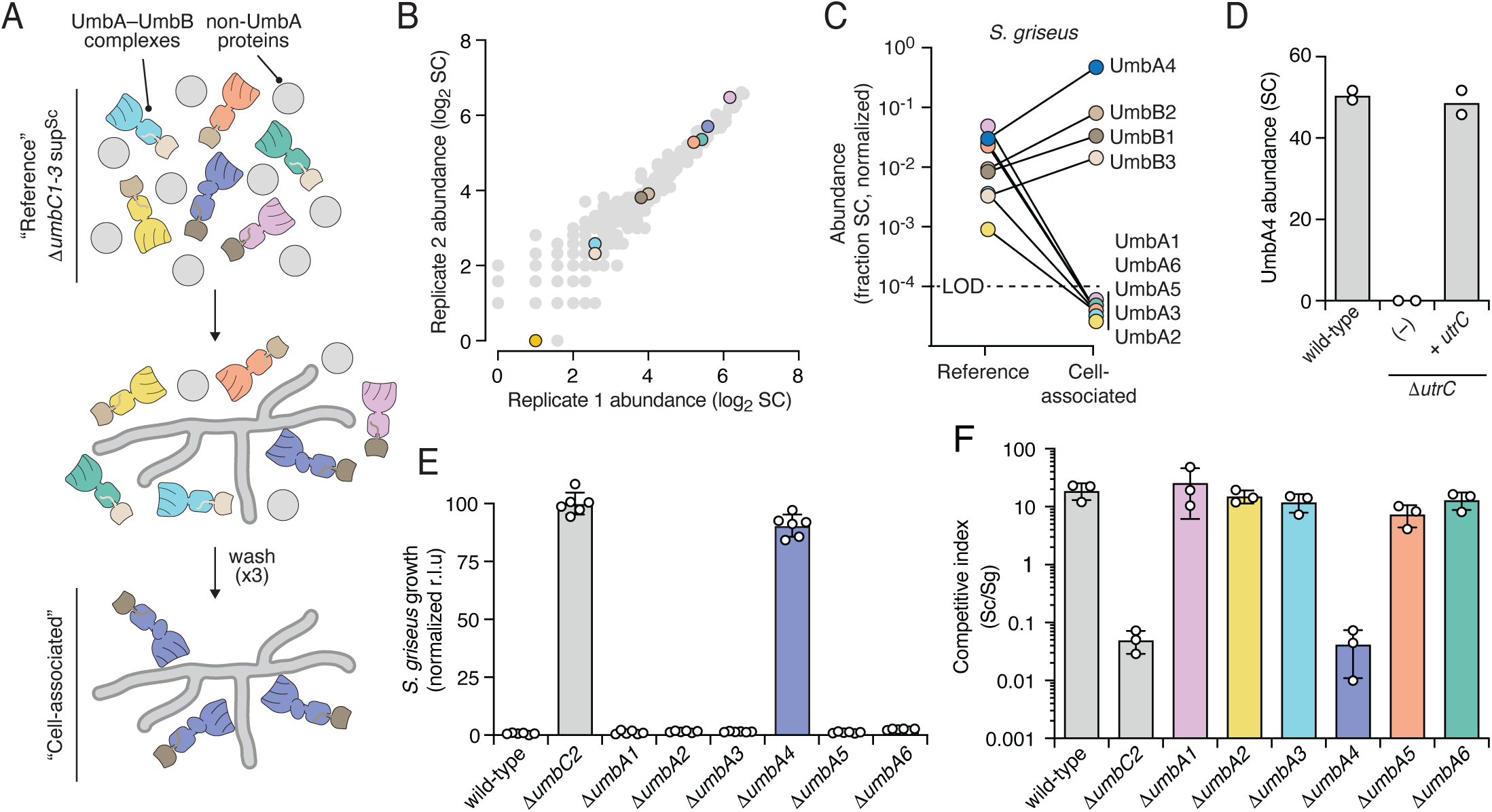
UmbA4 is required for targeting Umb2C of *S. coelicolor* to TUA–WTA producing *S. griseus.* A) Schematic representing the bacterial cell co-precipitation assay to identify *S. griseus*-interaction proteins in sup^Sc^. B) LC-MS-MS-based comparison of protein abundance across two independent replicate samples of *S. coelicolor ΔumbC1-3*-derived sup^Sc^ (“Reference” samples; SC, spectral counts). Spots representing UmbA and UmbB proteins are colored as shown in panel (C). C) Comparison of the proportion of spectral counts represented by UmbA and UmbB proteins in reference and *S. griseus* cell-associated samples of *S. coelicolor ΔumbC1-3*-derived sup^Sc^, identified as depicted in (A) (LOD, limit of detection,). Data represent means from two biological replicates. D) UmbA4 abundance following cell co-precipitation with the indicated strains of *S. griseus*. E) Growth yields of *S. griseus* (measured as relative luminescence units (r.l.u.)) after 16 h of treatment with sup^Sc^ derived from the indicated strains of *S. coelicolor*. Data represent the mean ± s.d. (*n* = 6). F) Outcome of growth competition assays between the indicated strains of *S. coelicolor* (Sc) and *S. griseus* (Sg).

We employed semi-quantitative mass spectrometry to determine the relative abundance of the UmbA proteins in our *S. coelicolor* Δ*umbC1-3*-derived sup^Sc^ samples. This method showed high reproducibility in the levels of the six *S. coelicolor* UmbA proteins across biological replicates samples (Figure 3B). Next we applied these samples to *S. griseus* target cells and evaluated the retention of individual UmbA proteins following extensive washing. False positives arising from peptide misidentification were ruled out by applying and analyzing in parallel a buffer-only control. Interestingly, among the UmbA proteins, only UmbA4 was detected in association with *S. griseus* (Figure 3C). As expected, based on the promiscuity in UmbB binding by orphan UmbA proteins, we also found UmbB1-3 enriched in the *S. griseus*-associated proteome. If TUAs are a receptor for the Umb2 particle of *S. coelicolor*, we reasoned that the interaction of UmbA4 with *S. griseus* should require their presence on the cell surface. Indeed, we found that UmbA4 is not retained on the surface of *S. griseus* cells lacking UtrC, and that UmbA4 interaction was restored by expression of UtrC from an ectopic locus (Figure 3D). These data show that UmbA4 interacts with *S. griseus* in a specific and TUA-dependent manner.

UmbA4 is an orphan UmbA present on all three Umb particles of *S. coelicolor*; however, the role of the protein in Umb2-based intoxication has not been measured. Moreover, the involvement of UmbA proteins generally in umbrella toxin function has not been determined. To explore these questions, we generated *S. coelicolor* strains bearing in-frame deletions in each of the *umbA* genes and applied sup^Sc^ derived from these to *S. griseus*. Remarkably, only inactivation of UmbA4 – the same UmbA we found interacts strongly with *S. griseus* cells – interfered with *S. griseus* intoxication. Sup^Sc^ derived from *S. coelicolor* Δ*umbA4* failed to strongly suppress *S. griseus* growth, similar to that of sup^Sc^ derived from *S. coelicolor* Δ*umbC2* (Figure 3E). In co-culture with *S. griseus*, secretion of active Umb2 particles confers substantial benefit to *S. coelicolor*. Using this assay, we also observed apparent inactivation of the Umb2 particle specifically by *umbA4* deletion (Figure 3F). Together, these data suggest that UmbA4 interaction with TUA molecules on the surface of *S. griseus* is required for Umb2-based intoxication.

### Two sites on the UmbA4 β-propeller recognize distinct TUA moieties

Our genetic and biochemical data suggest exquisite specificity of interaction between individual UmbA lectin domains and cell surface TUA oligosaccharides defines umbrella toxin targeting. To understand the molecular basis of this interaction, we sought to resolve the structure of a UmbA4–TUA complex using cryo-EM. Given the marked compositional heterogeneity of intact umbrella particles, and our inability to heterologously produce soluble UmbA4, we purified secreted UmbA4–UmbB1-3 complexes (UmbA4 associates with UmbB1-3) from *S. coelicolor* Δ*umbC1-3*. Cryo-EM analysis of these complexes revealed dispersed particles and enabled the determination of a 4.3 Å reconstruction of apo UmbA4 (Figure S3A-E and Table S4).

Addition of *S. griseus* TUA to UmbA4–UmbB1-3 complexes led to formation of helical filaments of ∼160 Å in width and >100 nm in length, as visualized by cryo-EM (Figure S4F-G). We therefore applied filament selection and subjected the extracted regions to helical refinement yielding a reconstruction at 3.3 Å resolution (Figure S3H-K). UmbA4 consists of three domains, an N-terminal trypsin-like domain, a central bulb lectin, and a C-terminal β-propeller (Figure 1A and Figure 4A-C), each of which were readily built into our maps. Formation of the helical filaments is mediated by the β-propeller domain and the resolution of the reconstruction gradually decreases towards the periphery of the helical assembly. As a result, the UmbB density which protrudes radially from the helix is more weakly resolved, which is further compounded by its compositional heterogeneity (Figure S3I). The map resolves TUA molecules bridging multiple UmbA4 β-propellers, explaining the oligosaccharide-induced oligomerization. The TUA was built into the cryo-EM density beginning at the non-reducing end, the location of which could be confidently deduced based on the termination of density and consideration of the steric restraints imposed by WTA attachment at the reducing end. Since the TUA polymers we purified are variable in length, we extended the TUA chain by cycles of repeating disaccharide addition and refinement until density no longer supported further addition. The final model comprises 12 monosaccharides: the unique non-reducing disaccharide and five repeating disaccharide units (Figure 4B and Table S4).

**Figure 4.**
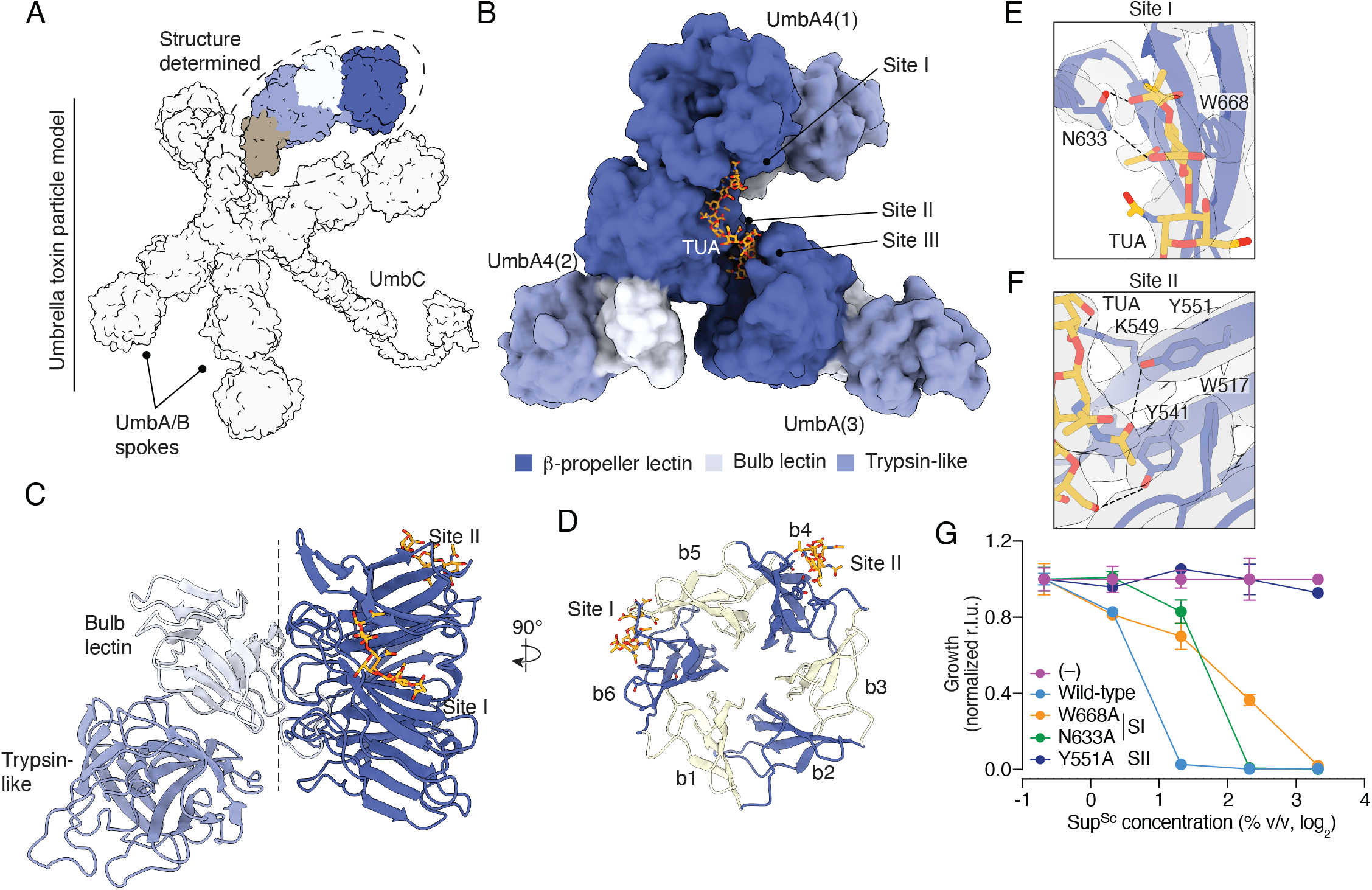
Structure of the UmbA4–TUA complex. A) Schematic depicting the *S. coelicolor* Umb1 particle indicating the localization of the UmbA–UmbB complex investigated here. B) Low pass filtered (6 Å) model derived from our cryoEM structure of three UmbA4 promoters bound to TUA purified from *S. griseus.* Three distinct sites on the β-propeller domains of different UmbA4 protomers that make contact with TUA are indicated. C, D) Side-on (C) and top-down (D) model of a single UmbA4 molecule indicating sites that make productive contact with TUA in different protomers in our cryoEM structure. Individual blades of the β-propeller are labeled in (D) (b1-6). E, F) Zoom in view of UmbA4–TUA interaction sites I (E) and II (F) indicating contacting amino acids. G) Growth yields of *S. griseus* following 16 hrs treatment with increasing concentrations of sup^Sc^ derived from *S. coelicolor* strains expressing UmbA4 variants with the indicated amino acid substitutions. UmbA4 levels were normalized across sup^Sc^ samples prior to dilution.

In our structure, a single TUA molecule contacts distinct sites (sites I-III) of the β-propeller domain within three UmbA4 protomers (Figure 4B). No contacts between the TUA and the trypsin-like and bulb lectin domains are present in our structure. Sites I and II occur within the canonical carbohydrate binding pockets of β-propeller lectins (Figure 4C,D). Site III, which we do not consider further, consists of superficial contacts between two UmbA protomers (Figure S4A,B). Site I, located between blades 5 and 6, accommodates the non-reducing end 4-OAc-ManNAc3NAcA-β(1→3)-GalNAc disaccharide and uronic acid moiety of the preceding repeat unit (Figure 4D,E). Site II, located on blade 4, interacts with an internal segment of TUA consisting of a repeating unit and a GalNAc residue from an adjacent repeat (Figure 4D,F). In an intact umbrella toxin particle, UmbA4 molecules are tethered to UmbC through interactions with UmbB, positioning them ∼100Å apart, which results in a distance between sites I and II on different UmbA4 molecules that exceeds the maximum length of TUA molecules. Thus, a single TUA chain is unlikely to bridge multiple UmbA4 molecules belonging to the same umbrella toxin particle on the target cell surface. It is conceivable that a long TUA molecule could span sites I and II on the same UmbA4; however, given energetic considerations and the size distribution of TUA we observed, we posit that most UmbA4 molecules interact with distinct TUA polymers at each binding site.

Like other β-propellers, the blades of UmbA4 are composed of a degenerate repeating sequence motif (Figure S4C-E)^25^. Sequences constituting the core fold of the blades and those forming the loops typically involved in Ca^2+^ binding (Ca^2+^ is not resolved in our structure) are highly conserved across blades, whereas the positions forming the carbohydrate binding pockets diverge substantially (Figure S4C). Nevertheless, our model suggests that residues occupying equivalent positions in sites I and II form a subset of the crucial contacts with TUA (Figure 4E,F). An example is Trp668 (Site I) and Tyr551 (Site II), which engage TUA through van der Waals interactions (Figure 4E,F). To test the functional significance of these interactions, we generated sup^Sc^ from *S. coelicolor* expressing UmbA4(W668A) or UmbA4(Y551A), normalized these samples to contain equivalent levels of UmbA4, and evaluated the capacity of each to intoxicate *S. griseus*. Remarkably, both single substitutions significantly impaired umbrella toxin activity, with UmbA4(Y551A) reducing it to the limit of detection (Figure 4G). These results motivated us to further probe predicted contacts in site I. Our model suggests Asn633 forms hydrogen bonds with both the carbonyl oxygen of 2-*N*-acetyl and the amide hydrogen of the 3-*N*-acetyl groups of the non-reducing TUA sugar (Figure 4E). Consistent with these interactions, we found sup^Sc^ derived from a strain expressing UmbA4(N633A) exhibits reduced capacity to inhibit *S. griseus* growth (Figure 4G). In total, our structural and physiological data offer a molecular model for the recruitment of umbrella toxin particles to the surface of target cells. They show that two positions within the β-propeller domain of UmbA4 engage in extensive contacts with distinct portions of the TUA polymer, offering an explanation for the fidelity of UmbA4-dependent recruitment of Umb2 particles to *S. griseus*.

### The Umb2 particle is recruited to target cells by multiple UmbA proteins

Our prior MS data suggests umbrella toxin particles, including Umb2, carry multiple UmbA proteins^10^. Taken together with our current findings, the model that emerges is that the different UmbA proteins associated with a given UmbC (via UmbB) could recruit the toxin to a multitude of target cells. Testing this model necessitated the identification of additional targets of the UmbC2 toxin of *S. coelicolor*. To this end, we screened a collection of diverse *Streptomyces* spp for susceptibility to sup^Sc^ (Figure S5A, Table S5). Subsequent experiments with one of the species hit in this screen, *S. achromogenes*, showed that its intoxication specifically requires UmbC2 (Figure S5B). With this UmbC2 target in hand, we proceeded to testing the strain for UmbA binding using our *S. coelicolor* Δ*umbC1-3* sup^Sc^ samples. Unlike *S. griseus*, which exclusively bound UmbA4, we found *S. achromogenes* specifically associated with UmbA5 (Figure 5A). Furthermore, Δ*umbA4*-derived sup^Sc^ intoxicated *S. achromogenes* indistinguishably from wild-type, and among all *umbA* deletion strains, only Δ*umbA5* lost the capacity to intoxicate *S. achromogenes* (Figure 5B). We conclude from these data that a single UmbC can be recruited to different target cells by different UmbA proteins.

**Figure 5.**
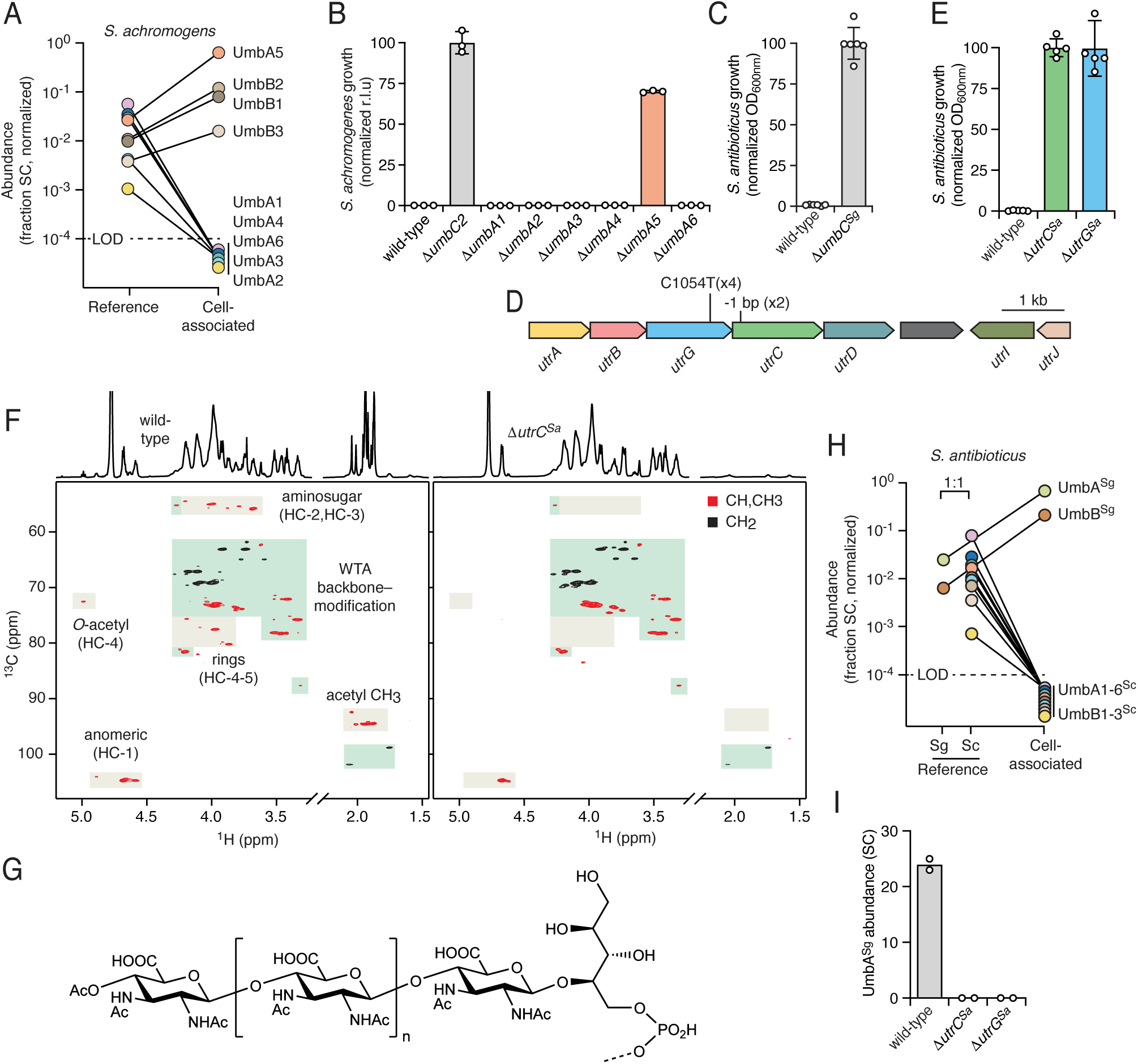
Divergent UmbA proteins mediate targeting of different species and recognize distinct TUA structures. A) Comparison of the proportion of spectral counts (SC) represented by UmbA and UmbB proteins in reference and cell-associated samples of *S. achromogenes* mixed with *S. coelicolor ΔumbC1-3*-derived sup^Sc^ (limit of detection, LOD). Data represent means from two biological replicates. See Figure 3A for experimental setup. B) Growth yields of *S. achromogenes* after 16 h of treatment with sup^Sc^ derived from the indicated strain of *S. coelicolor*. Data represent means ± s.d. (*n* = 3). C) Growth yields of *S. antibioticus* as measured by optical density (OD_600nm_) after 16 h of treatment with sup^Sg^ derived from the indicated strains of *S. griseus*. Data represent means ± s.d. (*n* = 6). D) Schematic indicating the location of mutations within the *utr* carbohydrate biosynthesis gene cluster of *S. antibioticus* selected during ALE with sup^Sg^ treatment. Numbers represent the number of times the indicated mutation was observed across six isolates sequenced. E) Growth of the indicated strains of *S. antibioticus* after 16 h of treatment with sup^Sg^. Data represent means ± s.d. (*n* = 5). F) Partial views of 2D multiplicity-edited ^1^H-^13^C HSQC NMR spectra of *S. antibioticus* wild-type (left) and Δ*utrC* (right) intact WTA polymers obtained by hydrolysis from isolated cell wall sacculi. The corresponding 1D ^1^H spectra are shown above each HSQC panel. Grey shading indicates peaks deriving from carbohydrates; green shading indicates poloyl*P*-derived peaks. G) Structure of the *S. antibioticus* TUA oligomer. The dashed bond indicates the expected linkage with the polyol-*P* WTA polymer. H) Comparison of the proportion of spectral counts (SC) represented by UmbA and UmbB proteins in reference and cell-associated samples derived from treatment of *S. antibioticus* cells with a 1:1 mixture of *S. coelicolor ΔumbC1-3*-derived sup^Sc^ and *S. griseus ΔumbC*-derived sup^Sg^ (limit of detection, LOD). For the reference samples, proteins derived from *S. griseus* and *S. coelicolor* are indicated below for clarity, and colors for *S. coelicolor* proteins correspond to panel (A). Data represent means from two biological replicates. G) UmbA^Sg^ abundance in cell-associated samples following precipitation from sup^Sg^ with *S. antibioticus* cells of the indicated genetic backgrounds.

### A distinct TUA structure mediates targeting by a divergent UmbA protein

To this point, our studies concerning umbrella toxin production, activity and target cell recognition have been limited to the particles produced by *S. coelicolor*. However, comprehensive bioinformatic analyses of *umb* loci suggest that hundreds of *Streptomyces* spp and bacteria distributed across multiple additional families of Actinobacteria might also employ umbrella toxins to mediate interbacterial antagonism. Though itself a target of *S. coelicolor* Umb2, *S. griseus* is one such bacterium. It possesses the minimal genetic elements we posited encode the necessary subunits of a minimal active umbrella particle: a single UmbA (UmbA^Sg^), UmbB (UmbB^Sg^), and a toxin-domain containing UmbC (UmbC^Sg^) (Figure S6A,B). Additionally, immediately downstream of *umbC^Sg^* is a gene encoding the predicted immunity factor for the toxin, UmbD^Sg^.

To probe the function of umbrella toxins outside of *S. coelicolor*, we generated an in-frame deletion in *S. griseus umbC* and compared the capacity of high molecular weight protein-enriched *S. griseus* supernatant (sup^Sg^) from this strain versus the wild-type to inhibit a panel of diverse *Streptomyces* spp. Similar to the behavior of sup^Sc^, we found that sup^Sg^ potently and selectively inhibits the growth of other *Streptomyces* spp in a *umbC*-dependent manner (Figure S6C, Table S5). Of the 37 strains included in this screen, we found one apparent target of the *S. grisues* umbrella toxin, *S. antibioticus*, a genetically tractable strain known for its production of actinomycin (Figure 5C)^26^. These findings provide initial evidence that umbrella toxins beyond those produced by *S. coelicolor* act in an antibacterial capacity.

Our identification of a second umbrella toxin producing organism and a cognate target allowed us to explore the generality of our findings pertaining to the significance of target cell surface carbohydrate recognition by UmbA proteins. We subjected *S. antibioticus* to selection in the presence of sup^Sg^, essentially as described above for *S. coelicolor* targets. After five passages, we screened 96 isolated clones and confirmed eight as highly resistant to sup^Sg^ (Figure S6D). Whole genome sequencing of six resistant isolates strongly implicated the *utr* gene cluster of *S. antibioticus* in Umb particle resistance. Non-synonymous mutations in *utrC* and a gene encoding a second glycosyltransferase in the TUA operon, *utrG*, were the only mutations we identified in multiple sequenced isolates (Table S6, Figure 5D). We generated strains containing in-frame deletions in each of these genes and confirmed that inactivation of either confers resistance to sup^Sg^ (Figure 5E).

The similarities between the TUA biosynthetic gene clusters of *S. griseus* and *S. antibioticus* are unambiguous; however, the two gene clusters also show substantial differences. For example, the *S. antibioticus* gene cluster lacks genes encoding two predicted epimerases and a glycosyltransferase found in *S. griseus* (*utrE, utrF* and *utrH*). We hypothesized that the differences in gene content between the TUA clusters of *S. griseus* and *S. antibioticus* are reflected in structural differences in their TUAs. This structural divergence would explain how *S. griseus* avoids umbrella particle adsorption to its own cell surface.

To test this hypothesis, we used NMR to determine the structure of the major species within material recovered from purified cell wall sacculi of *S. antibioticus* following TCA hydrolysis. NMR spectra indicated the presence of carbohydrate residues specific for TUA as well as polyol*P* residues from WTA (Figure 5F, Figure S7A,B and Table S7). DOSY analysis of the material revealed peaks attributable to TUA and WTA exhibit identical hydrodynamic properties, as observed in *S. griseus* (Figure S7E). Our analyses showed that the TUA polymer of *S. antibioticus* consists of β-1,4-linked GlcNAc3NAcA monosaccharide residues, rather than the distinct repeating disaccharide unit found in *S. griseus* (Figure 5F,G, Figure S7A,C, Table S7). We found acetylation at the 4-hydroxy position of the non-reducing end sugar is a common feature of the TUA structures. As in *S. griseus,* the reducing-end GlcNAc3NAcA in the TUA-like structure forms a β-1,4 glycosidic bond with a ribitol-5*P* residue. Also similar to *S. griseus*, we found that WTA portion of the TUA–WTA hybrid molecule of *S. antibioticus* is composed of ribitol phosphate, though in the latter the 1,5-linked ribitol*P* is heavily decorated with glucose, a modification not observed in *S. griseus* (Figure S7D). Additional *S. antibioticus* WTA substitutions included lysine and phosphoethanolamine. Purification and NMR analysis of equivalent material obtained from *S. antibioticus* Δ*utrC* revealed it contains similar WTA but with an additional unsubstituted terminal ribitol-5*P*, and it lacks TUA (Figure S7A,D, Table S7). These findings show that TUA–WTA hybrid molecules are found in Streptomyces beyond *S. griseus*. Furthermore, they are consistent with our hypothesis that differences in TUA structure between species determines umbrella toxin susceptibility.

Though the C-terminal lectin domain of UmbA^Sg^ adopts a predicted β-propeller fold, its amino acid sequence diverges significantly from the lectin domains of *S. coelicolor* UmbA1-6 (20-30% identity). If UmbA proteins define umbrella toxin particle target range by selective binding to variable cell surface carbohydrates as our data suggest, we reasoned that this sequence divergence could explain why the single umbrella particle elaborated by *S. griseus*, and none of the three produced by *S. coelicolor*, is able to target *S. antibioticus* (Figure S5A and Zhao et al^10^). To test this, we prepared a 1:1 mixture of sup^Sg^ and sup^Sc^, applied this to *S. antibioticus*, and used MS to evaluate the retention of Umb proteins after washing. Despite the multitude of UmbA proteins in sup^Sc^, we detected only UmbA^Sg^ and UmbB^Sg^ in association with *S. antibioticus* (Figure 5H). Consistent with our investigation of *S. griseus* targeting by *S. coelicolor*, UmbA^Sg^ binding to *S. antibioticus* required production of TUA (Figure 5I).

### Diversity of UmbA lectin domains and TUA biosynthetic gene clusters

Our finding that differences in TUA structure between *Streptomyces* dictate binding by specific UmbA proteins led us to hypothesize that an evolutionary arms race may be at play between toxin producing organisms and their would-be targets. This is consistent with our repeated observation that closely related *Streptomyces* species exhibit differential susceptibility to a given umbrella toxin particle. To investigate this hypothesis, we searched for evidence of predicted TUA structural diversity within the TUA biosynthetic loci of 15 species closely related to the umbrella-toxin target *S. griseus*. For reference, we also included a small group of more distantly related umbrella toxin target species and the model organism *S. coelicolor*. Using genomic comparisons and the results of our ALE experiments in *S. griseus* and *S. antibioticus*, we defined the bounds of the TUA biosynthetic cluster and used structural modeling and homology searching to predict the function of all intervening ORFs. Finally, we identified orthologous proteins by reciprocal best BLAST analysis. Consistent with umbrella toxins exerting selection pressure on TUA structures, our analysis revealed extensive variability in the genes present within TUA biosynthetic loci (Figure 6A). For example, the TUA gene cluster of *S. anulatus* lacks predicted epimerase and *O*-acetyl-transferase genes found in its close relative *S. griseus*. These differences offer an explanation for our finding that *S. anulatus* is not targeted by *S. coelicolor* umbrella toxins despite its overall high genomic similarity to *S. griseus* (Fast Average Nucleotide Identity of 92.4% across the genome^27^).

**Figure 6.**
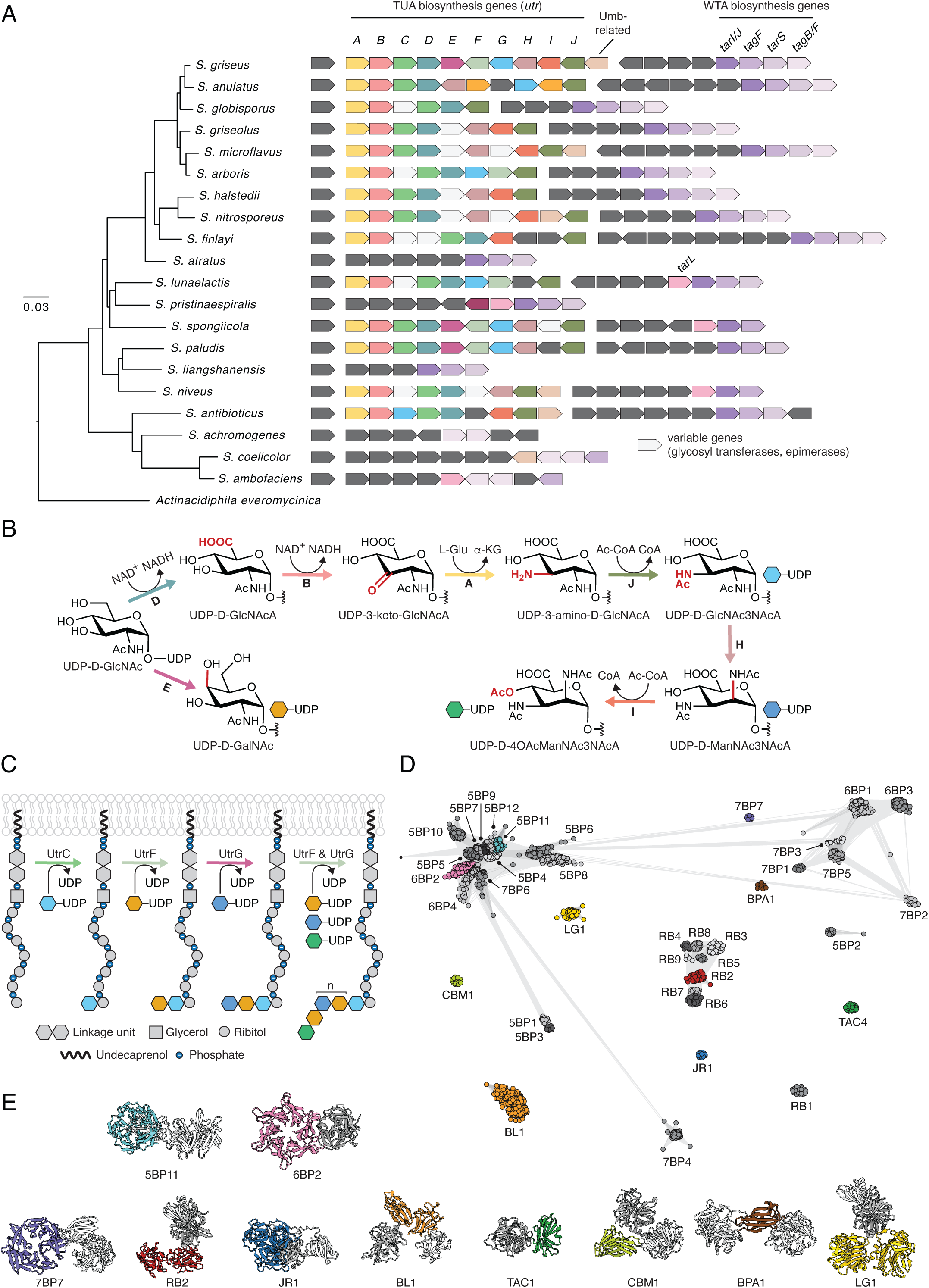
Diversity of *Streptomyces* TUA biosynthetic gene loci and UmbA lectin domains. A) Phylogeny (derived from the bacterial Genome Taxonomy Database (GTDB)) of selected *Streptomyces* species indicating gene content of TUA and linked WTA biosynthetic loci. Shared coloring indicates orthologous genes involved in TUA or WTA biosynthesis; all genes encoding umbrella toxin particle subunits are colored the same, regardless of orthology. TUA biosynthetic genes lacking orthologs in *S. griseus* are indicated in light grey and dark grey indicates genes with unrelated functions. Genes not drawn to scale. B, C) Predicted pathway for the biosynthesis of TUA by the Utr pathway of *S. griseus*. Key changes for each step in sugar precursor biosynthesis (B) are highlighted in red, and arrows representing the enzymes responsible for each reaction are colored according to panel (A). Colors of TUA monosaccharides and biosynthetic enzymes involved in polymer generation on WTA ribitol phosphate (C) correspond to panels (B) and (A), respectively. D) Network clustering analysis of UmbA c-terminal domains from diverse *Streptomyces* species. Only those domain families containing at least three family members and predicted to act as lectins are shown for clarity. (BP, b-propeller; RB, ricinB lectin; LG, laminin G; JR, jellyroll-like; BP, BPA-Ig; CB, CBM_4_9; BL, bulb lectin; TAC, TAC4. E) Structural models of UmbA proteins with C-terminal domains representative of different families. Labeled domains are colored according to panel (D). All other UmbA domains depicted in gray.

Functional predictions for the TUA genes of *S. griseus* and *S. antibioticus*, together with our elucidation of the corresponding chemical structures they produce, enabled us to generate a plausible TUA biosynthetic pathway (Figure 6B,C). It further permitted prediction of the extent to which TUA structure could vary across Streptomyces. In *S. griseus*, we found evidence that the TUA precursor UDP-ManNAc3NAcA is produced from UDP-GlcNAc by a similar pathway to that found in *Pseudomonas aeruginosa*, which incorporates ManNAc3NAcA into lipopolysaccharide^28^. Enzymes involved in this pathway include a UDP-sugar dehydrogenase (*utrD*), a sugar amino transferase and oxidase pair (*utrA, utrB*), an N-acetyltransferase (*utrJ*), and a UDP-glucose 2-epimerase (*utrH*). With the exception of *utrH*, we find that all of these genes are conserved across *Streptomyces* TUA biosynthesis gene clusters. This agrees with previous studies showing that the uronic acid component of TUA in these organisms consistently contains acetamido groups at the 2 and 3 positions^17^. The *S. griseus* TUA gene cluster additionally encodes a UDP-glucose 4-epimerase, *utrE*, which we predict is responsible for generating UDP-GalNAc, the second sugar of the repeating unit, from UDP-GlcNAc.

Beyond the precursor biosynthetic genes, only *utrC* is conserved throughout TUA loci (Figure 6A). UtrC bears closest structural resemblance to TarS, responsible for polyribitol phosphate β-O-GlcNAcylation in *S. aureus* (Figure S1C and Table S2)^13^, leading us to predict it initiates synthesis of TUA by linking GlcNAc3NAcA to ribitol phosphate. Supporting this, our genetic and chemical structural analyses indicate that the WTA of *S. griseus* Δ*utrC* bears no detectable TUA modifications (Figure 2A). In *S. griseus*, the next step, addition of α(1→4) linked GalNAc, is likely carried out by UtrF (Figure 6C). This enzyme is not found in *S. antibioticus*, consistent with a lack of α linkages in the TUA from this species (Figure 5G). We predict the repeated disaccharide portion of the TUA polymer of *S. griseus* is generated by alternating UtrG-catalyzed addition of β(1→3) linked ManNAc3NAcA and installation of α(1→4) linked GalNAc by UtrF. In *S. antibioticus,* UtrG would catalyze the addition of the repeated β(1→4) linked GlcNAc3NAcA residues that constitute its TUA polymer. We postulate that *O*-acetylation of the non-reducing end sugar in *S. griseus* and *S. antibioticus* is performed by UtrI. This function is patchily distributed across TUA-producing species, suggesting this terminal sugar modification varies. TUA gene clusters can encode additional glycosyltransferases and epimerases, indicative of further sources of TUA structural diversity.

A subset of the species we examined appear to lack TUA biosynthetic capacity entirely. Of note, we found that TUA genes are always linked to a gene encoding a putative bifunctional ribulose 5-phosphate reductase/CDP-ribitol pyrophosphorylase (*tarIJ*), which is required for ribitol WTA biosynthesis^29^. Species lacking this gene, and thus predicted to produce glycerol WTA, also lack TUA biosynthesis genes, suggesting that the linkage between TUA and ribitol WTA we uncovered may be a conserved feature of TUA across *Streptomyces.* Interestingly, species lacking TUA biosynthesis genes include two targets of *S. coelicolor* umbrella toxins, *S. ambofaciens* and *S. achromogens*, suggesting UmbA lectin domains may recognize additional cell surface carbohydrates beyond the hybrid TUA–WTA polymers identified here.

If an arms race scenario is mediating evolution of umbrella toxins and their targets, we would predict that UmbA lectin domains reflect a similar degree of diversification as their target molecules. To evaluate this, we identified UmbA-encoding genes among a collection of 11,223 Streptomycetaceae genomes, resulting in a total of 9859 UmbA sequences. Using domain scanning, network clustering analyses and structural predictions, we found the c-terminal domains of *Streptomyces* UmbA proteins represent 41 distinct families, encompassing 12 different protein folds (Figure 6D,E and Table S8). We found that the two UmbA proteins for which we identified a target molecule, UmbA4^Sc^ and UmbA^Sg^, contain 6-bladed β-propeller domains belonging to two distinct families (6BP-4 and 6BP-2, respectively). The divergence between these proteins is reflective of the structural differences between the TUA molecules they target. The UmbA required for targeting of *S. achromogens*, UmbA5^Sc^, contains a 5-bladed β-propeller domain. This species lacks the genes for producing TUA, suggesting UmbA5^Sc^ and related proteins likely recognize a distinct molecule. For the majority of the lectin families identified, target organisms and molecules remain uncharacterized. Given the sequence and protein fold diversity these families represent, we speculate that the surface carbohydrates exploited as receptors for umbrella toxins can vary widely. Additionally, we observe many instances in which a single UmbA protein contains more than one lectin domain, similar to UmbA4^Sc^. Our structural analysis indicated that the bulb lectin domain of UmbA4^Sc^ is not involved in *S. griseus* TUA recognition, and our genetic studies demonstrate that TUA targeting is necessary and sufficient for UmbA4^Sc^ recognition of *S. griseus*. This raises the possibility that UmbA proteins with multiple lectin domains recognize multiple, independent targets.

## Discussion

In this study, we present evidence that umbrella toxin targeting drives cell surface carbohydrate diversification in *Streptomyces.* We show that distinct TUA structures mediate targeting by divergent UmbA proteins, and we highlight extensive gene gain and loss among TUA biosynthetic loci of closely related species. This pattern suggests that horizontal exchange of TUA-modifying enzymes, and resulting alterations to TUA structure, promotes escape from umbrella toxin targeting among *Streptomyces* species. We also identified two species, *S. ambofaciens* and *S. achromogens*, which lack the genes for TUA biosynthesis, and yet are targets of one or more umbrella toxin particles produced by *S. coelicolor*. This indicates that additional surface molecules beyond TUA–WTA must be exploited by umbrella toxin as receptors.

Consistent with a wide range of carbohydrate structures serving as receptors, we show that multiple lectin families have been co-opted and diversified in UmbA proteins. Overall, the arms race dynamic we find underlies the evolution of umbrella toxin particles and their receptors provides compelling evidence of the pervasiveness and strength of the selection pressure imposed by umbrella toxin-mediated competition among *Streptomyces* species.

The diversity of lectin domains among UmbA proteins is compounded by the extensive variability in UmbC toxin domains. These include nearly one-hundred distinct toxin families, many of which are predicted to act via different biochemical mechanisms^10^. Modular coupling of variable receptor-targeting domains with divergent toxin domains in umbrella toxins is reminiscent of the contact-dependent growth inhibition (CDI) toxins of Proteobacteria. These toxins contain polymorphic cell recognition domains that interact with different outer-membrane proteins on target cells as well as a wide assortment of toxin domains^30,31^. Like umbrella toxins, CDI toxins exhibit narrow target specificity, typically mediating antagonism between strains of a given species. The convergent evolution of modularity among these toxins speaks to its selective benefit; whether this stems entirely from arms-race dynamics and the subversion of resistance in target cells is yet to be explored.

The structure we determined of UmbA4 in complex with TUA purified from *S. griseus* provides an unprecedented view of the contacts made between a β-propeller lectin and its natural ligand. This revealed just two of the six potential binding sites of UmbA4 make significant contact with TUA. While we cannot rule out that one or more of the four unoccupied sites might bind TUA in another context, none appear sterically occluded in our structure. Moreover, we find that disruption of either of the two sites we defined significantly reduces toxicity of the umbrella toxin particle toward *S. griseus*. These observations are consistent with the sequence divergence among UmbA4 blades and prior work suggesting that oligosaccharides approximating the physiological ligands of β-propellers are accomodated at fewer blade positions than are their simplified derivatives^25,32,33^. We speculate that limiting the binding of a particular TUA to a subset β-propeller blade positions in a given UmbA proteins could provide adaptive benefits. For instance, such an arrangement could broaden the species targeted by an umbrella particle by allowing a single β-propeller to recognize multiple ligands. Indeed, our finding that the two binding sites we identified interact with structurally distinct regions of TUA is consistent with this.

Our studies using intact cells to identify relevant UmbA proteins from Δ*umbC* strain-derived supernatant suggest that the individual lectins of these molecules interact tightly with their targets. Notably, β-propeller family lectins are those most prevalent in *Streptomyces* UmbA proteins, leading us to hypothesize that the avidity afforded by a single UmbA containing multiple carbohydrate binding sites, such as we identified on UmbA4, may be important for target cell recognition by umbrella toxins. Supporting this hypothesis, in UmbA proteins with other lectin folds, these domains frequently occur in multiple copies (Table S8). This avidity is likely amplified by the high local concentration of TUA–WTA at the binding interface on the cell surface.

If single UmbA proteins are sufficient to mediate target cell binding, why do umbrella toxin particles incorporate five UmbA-containing spokes? For species that encode more than one UmbA protein, this arrangement may allow targeting of more than one species or the subverting of resistance. When multiple UmbA proteins are encoded alongside multiple UmbC proteins, as in *S. coelicolor*, the five spoke arrangement could additionally serve as a bet hedging strategy that maximizes the likelihood of an effective lectin and toxin combination reaching a given target cell. Consistent with this, our work shows that UmbA4 and UmbA5 associate with all three Umb particles produced by *S. coelicolor*, yet only delivery of UmbC2 results in intoxication of *S. griseus* and *S. achromogens.* A non-mutually exclusive possibility is that while a single UmbA is sufficient for cell binding, interaction with a target cell surface that leads to intoxication may necessitate multiple points of contact. Future studies are needed to distinguish between these possibilities, and to characterize downstream steps in toxin translocation.

The TUA–WTA polymer we found can serve as an umbrella toxin receptor has not, to our knowledge, been previously observed. While the structure of TUA polymers in *Streptomyces* has received some attention^17^, the biological function of these molecules remains obscure. We readily obtained mutants lacking TUA biosynthetic capacity with no apparent growth defects *in vitro,* and our genomic analyses indicate TUA is not produced by all *Streptomyces* species, suggesting TUA production is dispensable for basic physiology. We posit that as a scaffold particularly amenable to modification, TUA may serve to enable escape from recognition by the many threats targeting surface-associated molecules. During selection with umbrella toxins derived from a single species, TUA deficient mutants have a distinct advantage, but natural populations are likely subject to simultaneous attack from many threats, including umbrella toxin particles from multiple species and phage, both of which could recognize the underlying WTA. Mutations conferring resistance to a particular threat may not be adaptive if they result in vulnerability to a different attack. Additionally, TUA may provide other benefits in the environment that are typically attributed to anionic cell wall polymers. In *Bacillus subtilis*, TUA is a component of the Pho regulon and is proposed to serve a role in conserving phosphate by substituting for WTA^34^. Given the linkage between WTA and TUA that we uncovered, it appears unlikely that TUA serves merely as a WTA equivalent under phosphate limitation in *Streptomyces.* Instead, TUA–WTA polymers could provide WTA-linked benefits such serving as a physical barrier to passage of harmful molecules, mediating attachment, or anchoring surface-associated proteins^35–38^.

Umbrella toxin particles and their individual components hold promise for a number of biotechnological applications. The vast diversity of lectin domains within UmbA proteins could be harnessed in situations where interaction with specific carbohydrates is of interest. Examples include detection of bacterial pathogens in food products, diagnosis of cancers based on alterations to cancer cell surface glycans, and generation of cancer biotherapeutics with lectin-based targeting moieties^39^. Intact or engineered umbrella toxin particles hold potential for development as antimicrobials. For instance, their modular nature could be exploited to generate chimeric particles for species-specific delivery of toxins of choice. Although to-date the only targets we have identified are *Streptomyces*, *umb* gene clusters are found across Actinobacteria^10^. This raises the possibility that umbrella toxin particles capable of intoxicating medically relevant Actintobacteria such as *Mycobacterium tuberculosis* and *Corynebacterium diptheriae* may await discovery. Advancing to targeting organisms beyond those naturally susceptible to umbrella toxins will likely require a mechanistic understanding of the steps downstream of receptor binding, an important avenue of future investigation.

## Acknowledgements

We thank David Brinkley and Yaxi Wang for assistance with figure preparation, and Simon Dove, Arne Rietsch, and members of the Mougous laboratory for helpful discussions. This work was supported by the U.S. Department of Energy, Office of Science, Basic Energy Sciences, Chemical Sciences, Geosciences and Biosciences Division, under award #DE-SC0015662, the Defense Advanced Research Projects Agency Biological Technologies Office Program: Harnessing Enzymatic Activity for Lifesaving Remedies (HEALR) under cooperative agreement no. HR0011-21-2-0012 (to J.D.M. and F.D.), the National Institute of Allergy and Infectious Diseases (DP1AI158186 and 75N93022C00036 to D.V.), an Investigators in the Pathogenesis of Infectious Disease Awards from the Burroughs Wellcome Fund (to D.V.), the University of Washington Arnold and Mabel Beckman cryo-EM center, the National Institutes of Health grant S10OD032290 (to D.V.) and the Saint louis University President’s Research Fund (to D.Z.).

J.D.M. and D.V. are HHMI Investigators, and D.V. and J.D.M hold the Hans Neurath Endowed Chair in Biochemistry and the Lynn M. and Michael D. Garvey Endowed Chair in Gastroenterology, respectively, at the University of Washington.

## Supplementary Figure Legends

**Figure S1.**
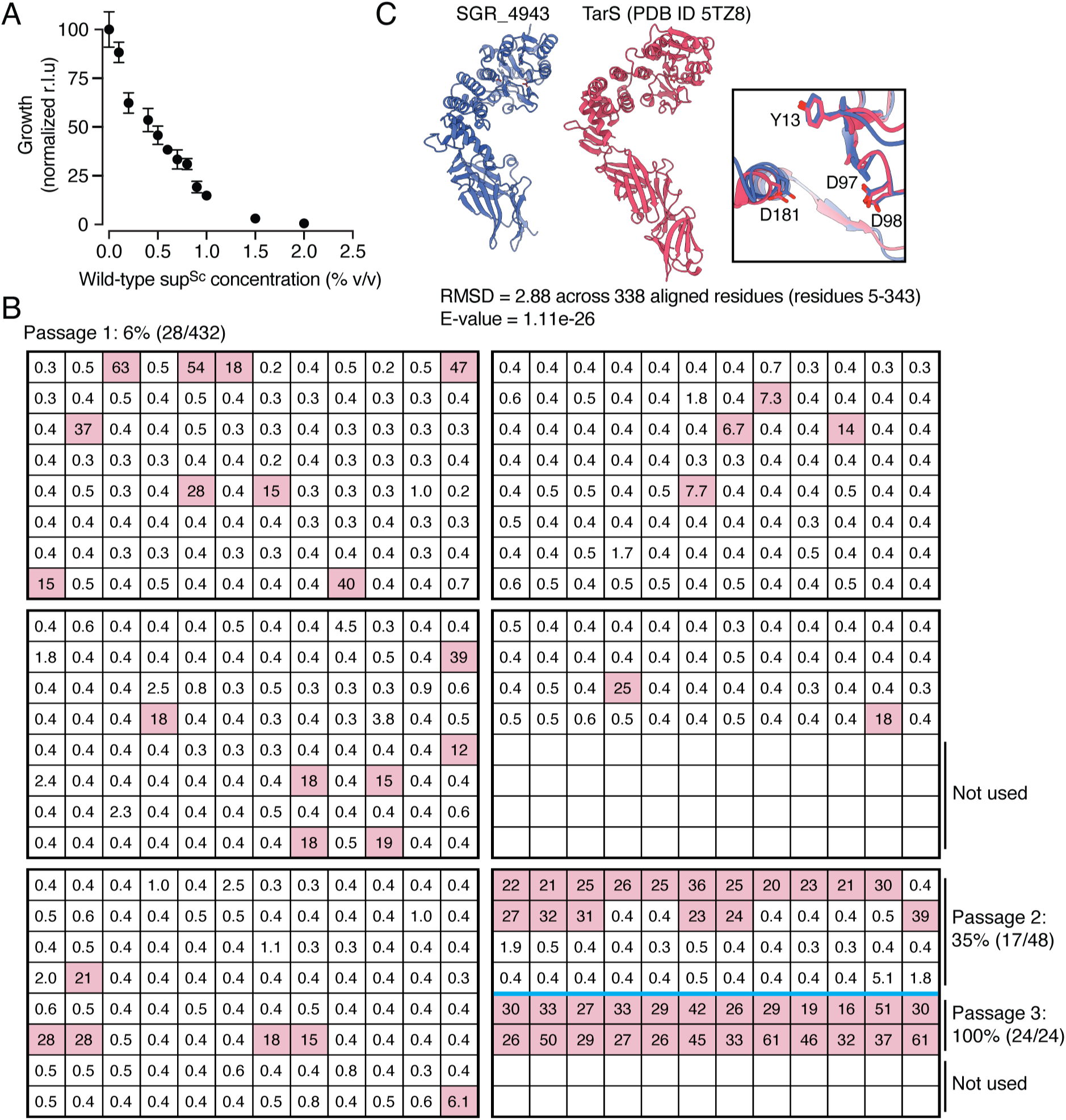
Isolation of *S. griseus* mutants resistant to sup^Sc^ via adaptive laboratory evolution. A) Growth of *S. griseus* after 16 h of treatment with wild-type sup^Sc^ serially diluted in *ΔumbC2* sup^Sc^. The concentration of wild-type sup^Sc^ in the treatment mixture is indicated. Data are normalized by the maximum and minimum levels of growth (measured as relative luminescence units (r.l.u.)), corresponding to treatment with only *ΔumbC2* or wild-type sup^Sc^, respectively. Data represent means ± s.d. (*n* = 3). B) Results of screen for sup^Sc^ resistance among evolved isolates from passages 1-3 of *S. griseus* ALE. Grids represent 96-well plates used to grow isolates for 16 h with sup^Sc^ treatment. Numbers indicate the ratio of evolved isolate growth to that of the wild-type ancestral strain. Isolates with a growth ratio exceeding 3 were considered resistant to sup^Sc^ and are illustrated by shaded wells. C) Predicted structures of the homologous regions of UtrC (SGR_4943) and TarS^14^. The magnified inset panel shows an overlay of the proteins’ predicted catalytic domain, including the catalytic residue D181.

**Figure S2.**
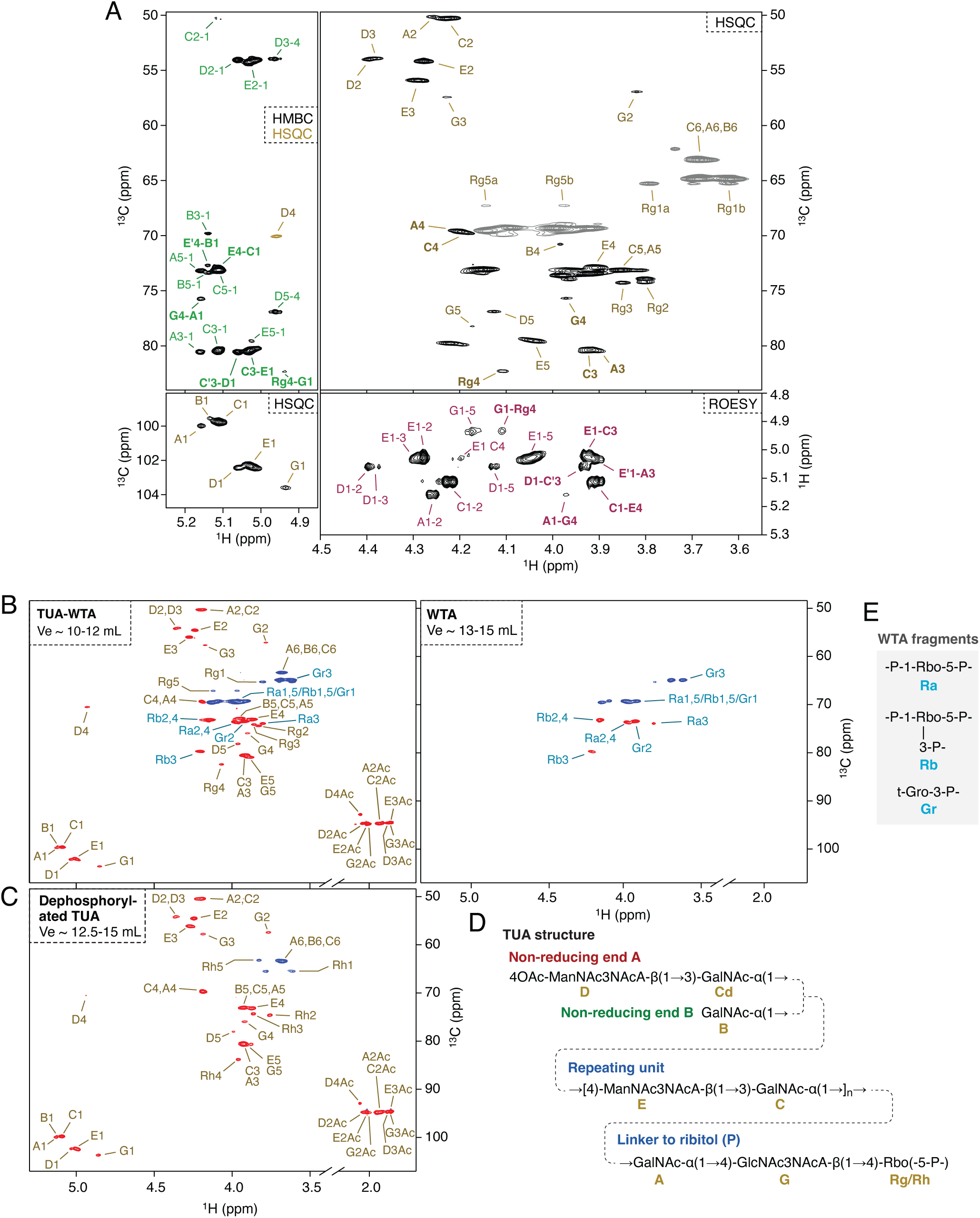
NMR-based structural analysis of WT *S. griseus* TUA-WTA. A) NMR structural analysis of TUA-WTA total TCA hydrolysate from *S. griseus*. Top right, 2D multiplicity-edited ^1^H,^13^C-HSQC NMR spectrum with carbohydrate signals labeled using residue codes from panel D. Signals of glycosidically linked positions are labeled in bold. For clarity, polyol-P signals of the WTA were not labeled. Positive signals (CH groups) are drawn in black and negative ones (CH2) in grey. Bottom right, a region of ^1^H,^1^H-ROESY spectrum that contains through-space correlation signals originating from the anomeric hydrogens, as well as H-4 of the acetylated residue D. Signal labels consist of the residue code and the ring position number of each of the two interacting nuclei. E.g., A1-2 marks signal due to correlation between H-1 and H-2 in residue A. Correlations between two residues across the glycosidic bond are labeled in bold (e.g., G1-Rg4). Bottom left, anomeric region of the ^1^H,^13^C-HSQC spectrum. Top left, an overlay of ^1^H,^13^C-HMBC (black signals) and ^1^H,^13^C-HSQC (olive signal) spectra in the ^13^C region up-field from the anomeric signals shown in the panel below. The HMBC signals show correlations between the anomeric hydrogen and carbon-2, -3 and/or -5 within a residue (e.g., A3-1). Further, the HMBC signals include inter-residue correlations through glycosidic bonds between anomeric hydrogen and the closest carbon in the aglycon (labeled in bold, e.g., G4-A1). B) 2D multiplicity-edited ^1^H,^13^C-HSQC NMR spectra of the larger TUA-WTA and the smaller WTA obtained from SEC of the total WT *S. griseus* cell wall TCA hydrolysate (Figure 2C). Signal assignments are based on the analysis of a full set of 2D NMR experiments acquired for each sample. Signals of TUA carbohydrate and the linking Rbo-5*P* are labeled in brown, while the signals of WTA polyol-*P* residues in blue. Positive signals (CH and CH_3_) are shown in red and negative signals (CH_2_) in blue. C) 2D multiplicity-edited ^1^H,^13^C-HSQC NMR spectrum of WT *S. griseus* dephosphorylated TUA obtained by SEC (Figure 2C). Signal assignments based on the analysis of a full set of 2D NMR experiments are indicated. D) Schematic structure of the TUA with NMR assignment code indicated in brown for each residue. The codes are as in panels A-C, and in Table S3. E) Major polyol-*P* units identified in the WTA, with NMR assignment codes indicated in blue.

**Figure S3.**
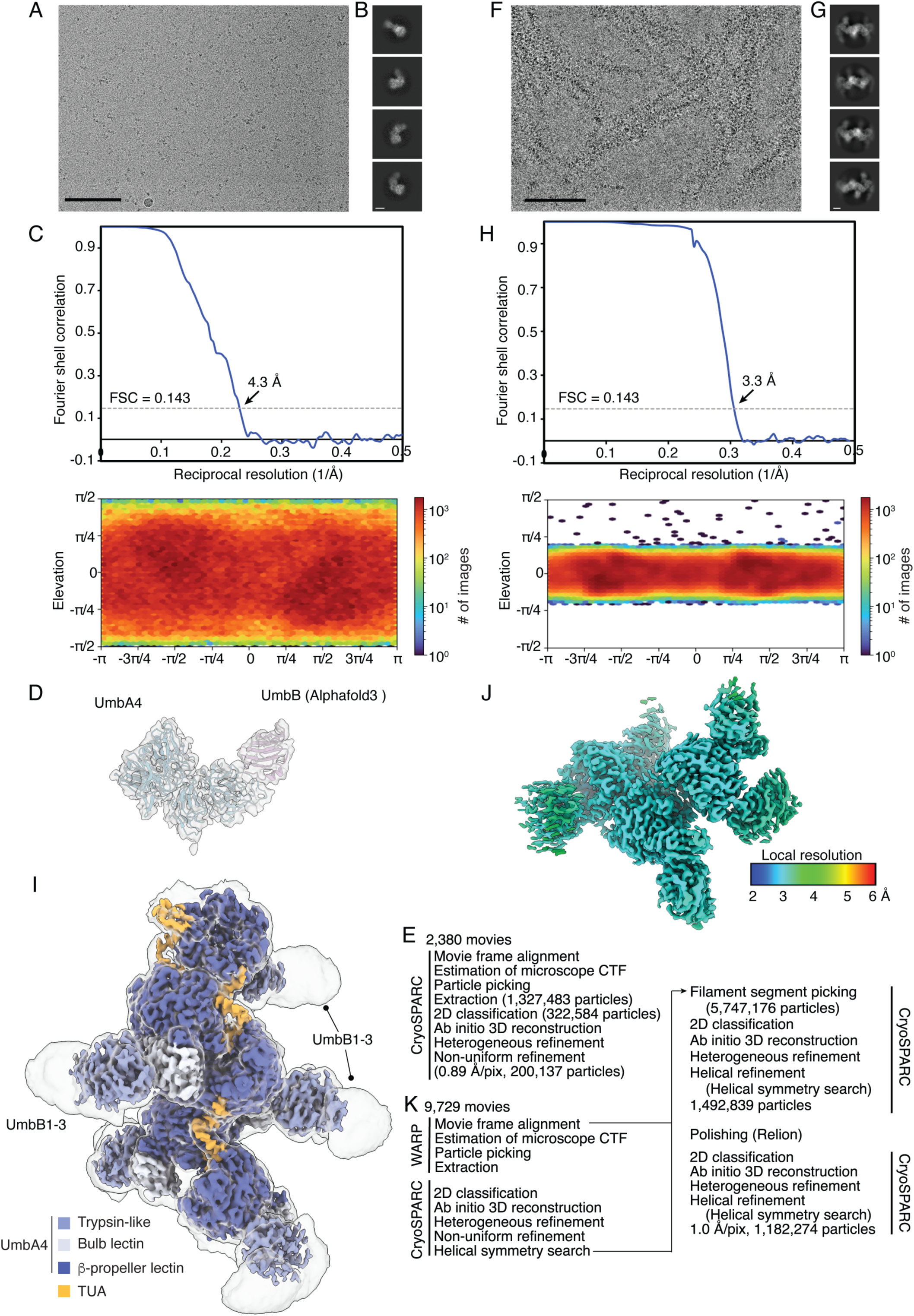
Cryo-EM data processing of the UmbA4 and TUA molecules bound UmbA4 complex datasets. A, B, F and G) Representative electron micrographs (A, F) and 2D class averages (B, G) of the UmbA4 complex (A, B) or TUA molecules bound UmbA4 complex (F, G) embedded in vitreous ice. Scale bars: 100 nm (A,F), 20 nm (B,G). C,H) Gold-standard Fourier shell correlation curve of the UmbA4 complex (C) or TUA molecules bound UmbA4 complex (H). The 0.143 cutoff is indicated by a horizontal dashed line. The angular distribution of particle images calculated using cryoSPARC is shown as a heat map below. D) 3D reconstruction of UmbA4 complex that encompassed two proteins (UmbA4 in blue and UmbB in magenta). E,K) Data processing flowchart. CTF: contrast transfer function; NUR: non-uniform refinement. J) Local resolution estimation of TUA molecules bound UmbA4 complex reconstruction calculated using cryoSPARC and plotted on the sharpened maps. I) Unsharpened (transparent) and sharpened (opaque) cryo-EM maps of a helical segment of the TUA-UmbA4-UmbB1-3 complex. Regions encompassing TUA and UmbA4 are colored based on model proximity.

**Figure S4.**
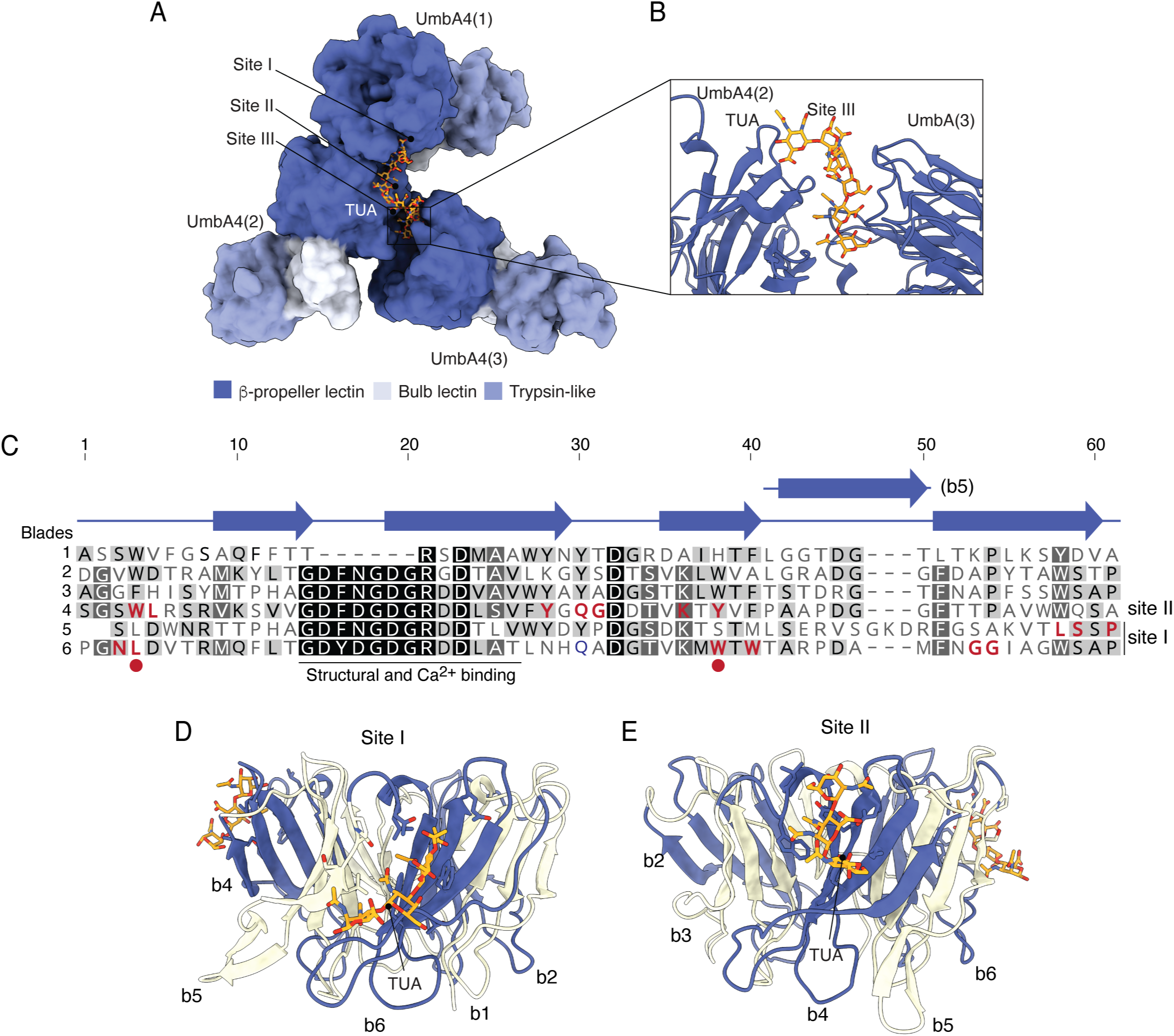
TUA binds at conserved sites on the b-propeller lectin. A) Overview of sites I, II, and III on the assymetric unit containing three UmbA4 protomers. B) Close-up of site III. TUA packs between UmbA4(2) and UmbA(3) making weak contacts with both protomers. C) Sequence alignment of the b-propeller lectin blades. Secondary structure is indicated above the alignment. Residues that form direct interactions with TUA are indicated (red). Residue positions that interact with TUA in both site I and II are indicated (circle). Positions of TUA in D) site I, between blades 5 and 6 (B5, B6) and in E) site II, within blade 4 (B4).

**Figure S5.**
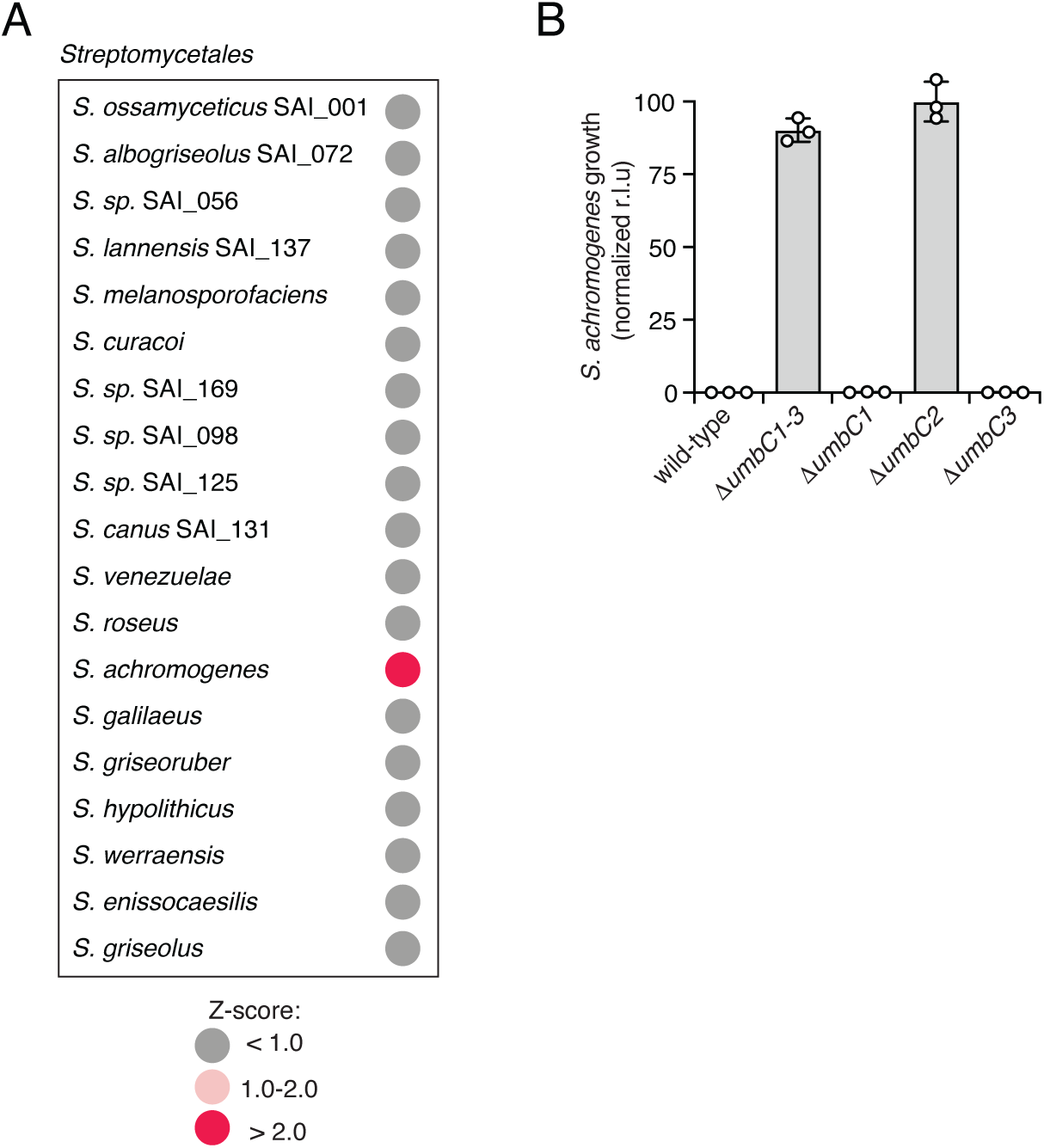
*S. achromogenes* is susceptible intoxication by UmbC2 of *S. coelicolor*. A) *S. coelicolor* Umb toxin susceptibility screening results. *Z* scores calculated from ratio of growth in control supernatant to growth in sup^Sc^ from two biological replicates of the screen; scores >2 indicate significant Umb-dependent inhibition. Raw data provided in Table S5. B) Growth yields of *S. achromogenes* (measured as relative luminescence units (r.l.u.)) after 16 h of treatment with sup^Sc^ derived from the indicated strain of *S. coelicolor*. Data represent the mean ± s.d. (*n* = 3).

**Figure S6.**
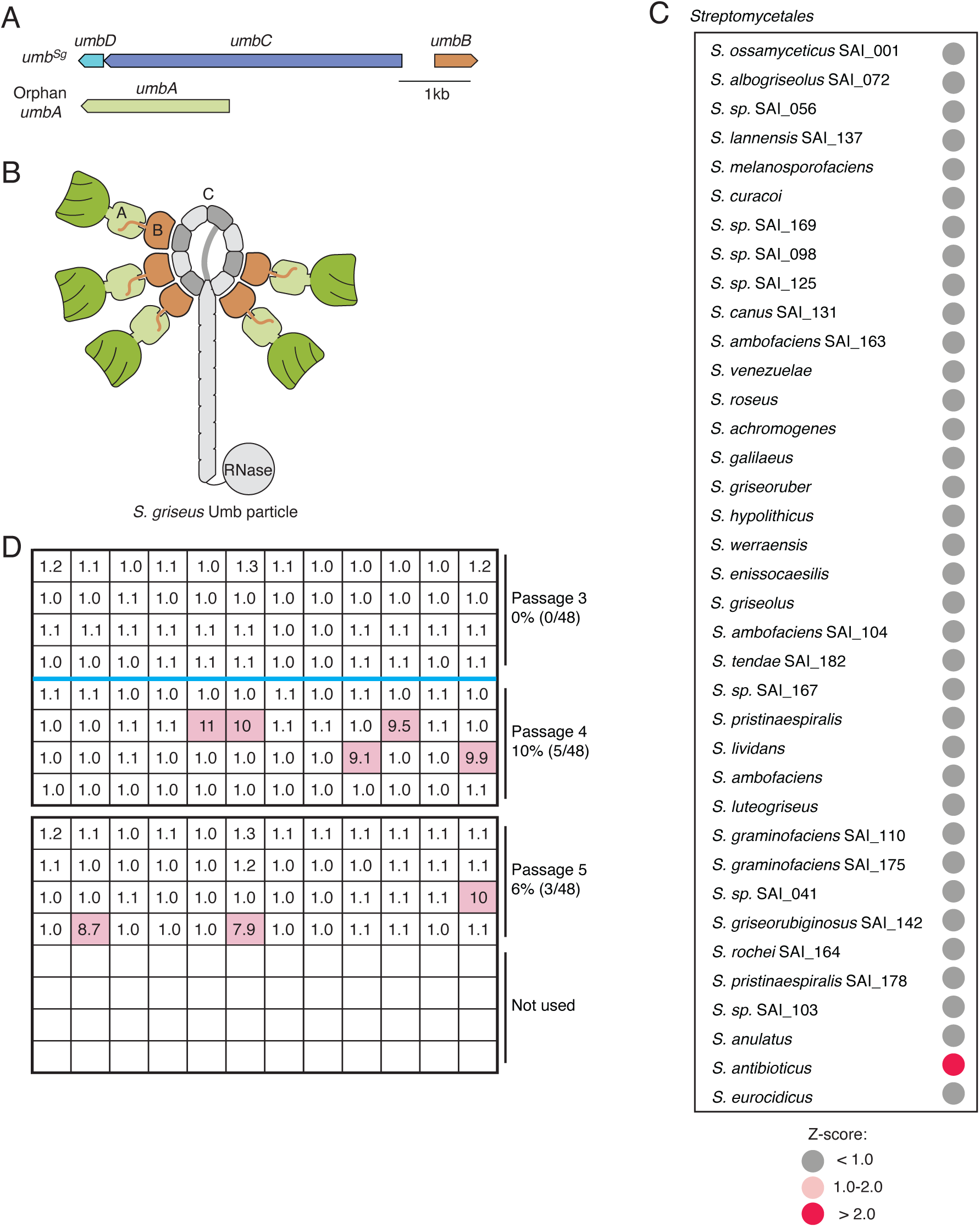
Identification and ALE of a target of *S. griseus* umbrella toxin particle. A) Loci encoding Umb protein complex components in S. griseus. UmbA is encoded distantly from other complex components. B) Schematic illustrating the composition of the single umbrella toxin particle produced by *S. griseus.* Toxin domain labeled according to predicted RNase activity. C) *S. griseus* Umb toxin susceptibility screening results. *Z* scores calculated from ratio of growth in control supernatant to growth in sup^Sg^ from two biological replicates of the screen; scores >2 indicate significant Umb-dependent inhibition. Raw data provided in Table S5. D) Results of screen for sup^Sg^ resistance among evolved isolates from passages 3-5 of *S. antibioticus* ALE. Grids represent 96-well plates used to grow isolates for 16 h with sup^Sg^ treatment. Numbers indicate the ratio of evolved isolate growth to that of the wild-type ancestral strain. Isolates with a growth ratio exceeding 3 were considered resistant to sup^Sg^ and are illustrated by shaded wells.

**Figure S7.**
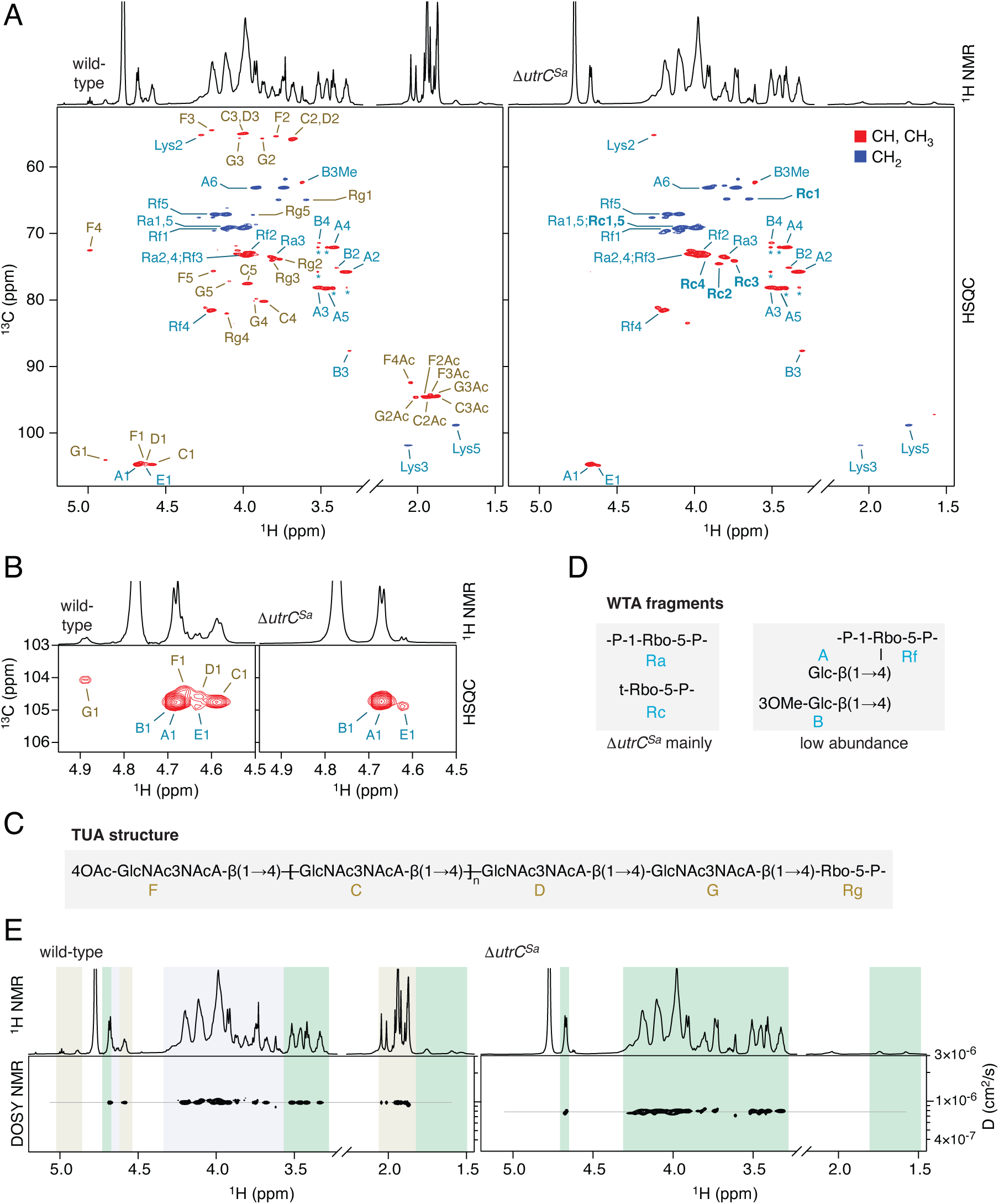
Structural analysis of *S. antibioticus* cell wall polymers. A) 2D multiplicity-edited ^1^H,^13^C-HSQC NMR spectra of *S. antibioticus* WT (left) and Δ*utrC* (right) intact polymers isolated from cell walls by acid hydrolysis. The corresponding 1D ^1^H spectra are shown above each HSQC panel. Signals of the TUA are labeled in brown and signals of WTA in blue. The labels show the residue code pertinent to the TUA and WTA structures in panels C and D, respectively, and C–H group number in that residue. Positive signals (CH and CH_3_) are shown in red and negative signals (CH_2_) in blue. Asterisks mark weak COSY-type peaks that formed between the strong signals of residue A. B) Expanded anomeric region of the same HSQC spectra shown in panel A. Signal and spectra colors have the same meaning as in panel A. C) Structure of the TUA oligomer with a repeating unit consisting of a single →4)-GlcNAc3NAcA-β(1→ residue. The TUA is terminated with a 4O-acetylated GlcNAc3NAcA residue on the non-reducing end, while the reducing-end residue of GlcNAc3NAcA forms a β(1→4) glycosidic bond with a ribitol-5-phosphate residue. D) Ribitol phosphate units identified in the WT and mutant WTA. The WTA contains 1,5-linked ribitol-*P* residues, a large portion of which is substituted by β-Glc residue in the 2- or 4-position. The mutant WTA has much higher content of a terminal ribitol-5*P* residues (residue Rc, in bold), presumably equivalent to the terminal Rbo-5P that is substituted by TUA in the WT material. E) ^1^H NMR (top) and DOSY (bottom) spectra of the WT (left) and Δ*utrC* (right) TUA/WTA preparations. The WT DOSY spectrum shows that the TUA and WTA have the same diffusion coefficient, indicating the same size. All WTA signals in the Δ*utrC* DOSY are also aligned at the same *D* level, consistent with a size-uniform preparation. Give its lower *D*, the size of Δ*utrC* WTA is likely somewhat larger compared to the WT TUA-WTA that exhibits a larger *D*. Gray, green and blue backgrounds mark regions with TUA only, WTA only and overlapping TUA+WTA signals, respectively.

## Methods

### Strains, media and growth conditions

Strains, plasmids and primers employed in this study are listed in Table S5 (*Streptomyces* species screened for susceptibility to umbrella toxins) and Table S9 (all other strains, plasmids and primers)*. Escherichia coli* strain DH5α was used for plasmid maintenance and ET12567(pUZ8002) for interspecies conjugation. *E. coli* strains were grown in Lysogeny broth (LB) at 37 ℃ with shaking at 200 r.p.m or on LB agar plates (2 %, w/v). Strains of *S. coelicolor* A3(2), *S. griseus* (NRRL B-2682), *S. antibioticus* (NRRL B-1701), and *S. achromogenes* (SANT-13) were used for studying Umb particles. Unless otherwise noted, *Streptomyces* strains were cultivated in R5, TSBY, or 2xYT liquid media in baffled flasks with 3 mm glass beads, shaken at 28 ℃ and 220 r.p.m., or on ISP2, ISP4 or SFM agar plates (2 %, w/v). Media were supplemented as needed with antibiotics, including apramycin, kanamycin, chloramphenicol, and trimethoprim (all used at 50 mg/L).

### Plasmid construction

Primers and synthetic DNA fragments used in this study were obtained from Integrated DNA Technologies and are listed in Table S9. Plasmid constructs were designed using Geneious Prime, generated via Gibson assembly, and confirmed by sequencing. *S. coelicolor* genetic manipulation was conducted using pKGLP2a with the apramycin resistance gene (*aac*(3)*IV*), as previously described^10^. To enhance the efficiency of *S. griseus* and S*. antibioticus* mutant generation, the *codA(sm)* counterselection marker and its promoter, amplified from pLQ752^40^, were introduced to pKGLP2a to generate pKGLP2a-*codA(sm)*. CodA, a cytosine deaminase, allows for counterselection via the addition of 5-fluorocytosine, which it converts into highly toxic 5-fluorouracil^41^. Constructs for introducing deletions in chromosome of *Streptomyces* strains using pKGLP2a or pKGLP2a-*codA(sm)* were generated through Gibson assembly with 1.5-2 kb arms flanking the modification site. The integrative vector pSET152, which directs integration at the *attB* phage attachment site, was used for gene expression and complementation in *Streptomyces*^42^. For expression of UmbA4 in *S. coelicolor* strains, the gene was fused sequentially at the C-terminus with a 2xGGGGS linker, a 3xK tag and a 6xH tag and at N-terminus with the *S. coelicolor* SCO4296 promoter (198 bp) by PCR, then inserted to pSET152 by Gibson assembly. UmbA4 variants with mutations in key residues involved in TUA interaction were generated by PCR or DNA synthesis. For *utrC* complementation in *S. griseus* Δ*utrC*, the gene was fused with the SCO4296 promoter at and cloned into pSET152.

### Genetic manipulation of *Streptomyces* strains

To generate genetically modified *S. coelicolor*, vectors derived from pKGLP2a with the corresponding inserts were introduced via *E. coli*-*Streptomyces* intergeneric conjugation using *E. coli* ET12567(pUZ8002) as the donor strain. Conjugation and mutant screening were performed as described previously^10^. For gene deletion in *S. griseus* and *S. antibioticus*, derivatives of pKGLP2a-*codA(sm)* carrying homologous alleles were transferred via the same conjugation method as for *S. coelicolor*, with an extended 50℃ heat-shock treatment of spores (20 min) before mixing with donor *E. coli* cells. Transconjugants were streaked onto ISP4 agar supplemented with trimethoprim and apramycin and then cultured in non-selective TSBY medium for 24 h. The cultures were subsequently streaked on non-selective ISP4 agar and incubated at 30 ℃ for 5-7 days to promote sporulation. Spores were collected, diluted, and plated on ISP4 agar containing 50 mg/L 5-fluorocytosine. After 2 days of incubation, colonies were screened for apramycin sensitivity and the presence of the desired allele via PCR. For gene expression or complementation in *Streptomyces* strains, the integrative vector pSET152 carrying the desired genes and corresponding promoters was introduced through conjugation, following the procedure species-specific procedures described above for pKGLP2a and pKGLP2a-*codA(sm)*.

### Preparation of concentrated supernatant from *Streptomyces* strains

Cultures of *S. coelicolor* used for concentrated supernatant preparation were grown as previously described^10^. For preparation of *S. griseus* concentrated supernatant (sup^Sg^), spores were grown in 30 mL R5 medium for 24 h, then back diluted to an OD_600_ of 0.005 in 50 mL R5 medium per flask, with a total combined volume of 150 mL. Back-diluted cultures were incubated for approximate16 h until OD_600_ reached 4-5. For both species, cultures were centrifuged to remove cells, resulting supernatant was filtered with a 0.45 μm PES membrane vacuum filter and then concentrated using 100 kDa cutoff concentrators (Amicon) until reaching a final volume of 3 mL. Concentrated supernatants were desalted using Econo-Pac 10DG desalting columns (Bio-Rad), filter sterilized (0.22 μm), aliquoted and stored at -80 ℃.

### Isolation and identification of Streptomyces strains from soil and root samples

We grew *Arabidopsis thaliana* (Col-0) under standardized conditions in natural soils from diverse U.S. regions. After 8 weeks, we collected bulk soil and roots for bacterial culturing. To enrich for endophytes, roots were vortexed and sonicated in sterile water, then homogenized and serially diluted (1/10) for plating on ISP4 and water agar; bulk soil was treated similarly. *Streptomyces*-like colonies were re-streaked twice on ISP4 to obtain pure cultures. Spores were collected in 40% glycerol and stored at -80 °C. To identify isolated *Streptomyces* species, genomic DNA was extracted from 20 h cultures using InstaGene Matrix (Bio-Rad), and the V1-V3 region of the 16S ribosomal RNA gene was amplified by PCR and analyzed by Sanger sequencing (Azenta). Species identification was performed using a BLAST search of these sequences against the 16S ribosomal RNA sequences (Bacteria and Archaea) database.

### Screening *Streptomyces* strains for sensitivity to Umb toxins

To screen for new target strains sensitive to *S. coelicolor* Umb particles, 19 diverse *Streptomyces* species (see Table S5 for complete list) were streaked on ISP4 and grown for 24-72 h before being inoculated into 100 μL TSBY cultures in a 96-well plate. The plate was incubated at 30℃ for 36 h in a BioTek LogPhase 600 Microbiology Reader with shaking at 800 r.p.m. Optical densities of the initial cultures were measured and diluted to an OD_600_ of 0.001 in TSBY medium. Next, 90 μL aliquots of each test strain culture were mixed with 10 μL of sup^Sc^ derived from either wild-type or Δ*umbC1-3* (negative control) in a 96-well plate. The plate was incubated at 30℃ for 16 h. Cell viability of the cultures was assessed by combining each with 100 μl BacTiter-Glo reagent (Promega), incubating for 5 min at room temperature, and measuring luminescence using a BioTek Cytation 1 imaging reader. To identify a strain sensitive to the *S. griseus* Umb particle, 37 diverse *Streptomyces* species (listed in Table S5) were screened using the same procedure described above, with minor modifications. In brief, 99 μL diluted culture aliquots were mixed with 1 μL sup^Sg^ from either wild-type or Δ*umbC S. griseus* (negative control). Cell viability was determined using the BacTiter-Glo reagent after incubation for 16 h. For both screens, growth inhibition was determined by calculating the ratio of luminescence signals from cultures treated with Δ*umbC* supernatant to those treated with wild-type supernatant. Two biological replicates were performed for each screen, and Z scores were calculated from the average of log_2_-transformed ratios.

### Evaluation of Umb particle toxicity

Methods of assessing the toxicity of Umb particles using concentrated supernatants from *S. coelicolor* wild-type and mutant strains (sup^Sc^) and *Streptomyces* co-culture competition assays have been described previously^10^. To evaluate the toxicity of Umb particles produced by *S. griseus,* concentrated supernatants from *S. griseus* wild-type and mutant strains (sup^Sg^ ) were prepared as described above. *S. antibioticus* precultures (grown for 20 h, as described above) were diluted to an OD_600_ of 0.01 in TSBY medium. A 99-µL aliquot of the diluted culture was mixed with 1 µL of sup^Sg^ in a 96-well plate and then incubated in a LogPhase for 16 h. Growth of *S. antibioticus* was quantified by OD_600_ measurement after 16 h of incubation.

### Adaptive laboratory evolution of Umb-resistant strains

To establish clonal ancestor populations for these experiments, wild-type *S. griseus* and *S. antibioticus* strains were streaked onto ISP4 agar and incubated at 30 ℃ for 5-7 days until isolated colonies sporulated. Spores from a single colony were collected in 20% glycerol (w/v) using a sterilized cotton swab, streaked onto multiple ISP4 agar plates, and incubated until sporulation. The spores were then collected in 20% glycerol (w/v), aliquoted, and stored at -80℃.

To determine the minimum inhibitory concentration (MIC) of sup^Sc^ toward *S. griseus*, sup^Sc^ was serially diluted 2-fold in sup^Sc^ from *S. coelicolor* Δ*umbC2* to create working solutions with different concentrations of Umb2 particles. *S. griseus* clonal spores were cultured in 30 mL TSBY medium for 20 h, then back diluted to an OD_600_ of 0.01. A 90 μL aliquot of the diluted culture was mixed with 10 μL of each working solution in a 96-well plate. The plate was incubated in a LogPhase at 30 ℃ for 16 h, and cell viability was assessed using BacTiter-Glo reagent. Data were normalized to the range of growth observed in positive (undiluted wild-type sup^Sc^) and negative (Δ*umbC2* sup^Sc^) control treatments. A similar procedure was used to determine the MIC of sup^Sg^ against *S. antibioticus*, except cell growth was assessed by OD_600_ measurement. The concentrations of sup^Sc^ and sup^Sg^ that resulted in approximately 90% growth inhibition were selected as the initial treatment for evolution of Umb-resistant mutants (1.25% for sup^Sc^ and 0.12% for sup^Sg^).

For ALE of *S. griseus* mutants resistant to *S. coelicolor* Umb2 particles, *S. griseus* clonal spores were grown in TSBY medium for 20 h then diluted to an OD_600_ of 0.01. 5 mL aliquots of diluted culture were added to 100 mL baffled flasks containing 3 mm glass beads and mixed with sup^Sc^ from wild-type or *S. coelicolor* Δ*umbC2* mutant to a final concentration of 1.25% (v/v) (passage 1). Three independent lineages were treated with wild-type sup^Sc^, while one lineage treated with *S. coelicolor* Δ*umbC2* sup^Sc^ served as a negative control. After 48 h of incubation, cultures reached an OD_600_ of 8-9 and were reinoculated into fresh TSBY medium containing 1.25% sup^Sc^ to an OD_600_ of 0.01 to initiate passage 2. Meanwhile, 50 µL cells from passage 1 were streaked on ISP4 agar plates for sporulation. The liquid cultures reached OD_600_ of 2-7 after 24 h of incubation, and were streaked out again and rediluted to OD_600_ of 0.01 in TSBY medium containing 1.25% sup^Sc^ for passage 3. In this passage, the OD_600_ reached 8-10 within 24 h of incubation, and cultures were streaked for sporulation at this point.

Spores from different lineages and passages were collected, diluted, and plated on ISP4 agar for single colony isolation. Colonies were selected for sup^Sc^ sensitivity testing at random in equal numbers from each lineage for a total of 432 colonies from passage 1, 48 colonies from passage 2, and 24 colonies from passage 3. Along with the ancestor strain, these individual colonies were used to inoculate 100 μL TSBY cultures in 96-well plates and incubated for 16 h at 30℃ in a BioTek LogPhase 600 Microbiology Reader. Optical densities of starter cultures were measured and used to inoculate 50 μL TSBY with 10% wild-type sup^Sc^ to an OD_600_ of 0.003 in new 96-well plates. These plates were incubated in a LogPhase for 16 h and growth was assessed by combining each with 50 μL BacTiter-Glo reagent and measuring luminescence after a 5-min incubation at room temperature.

*S. antibioticus* mutants resistant to *S. griseus* Umb particles were generated using a similar ALE procedure as described above, with the modifications noted below. In brief, pre-cultured *S. antibioticus* cells were diluted to an OD_600_ of 0.01 and treated with 0.12% (v/v) sup^Sg^ in passage 1. After 68 h of incubation, cultures reached an OD_600_ of 1.3-5.2 and were reinoculated into fresh TSBY medium containing 0.12% sup^Sg^ at an OD_600_ of 0.01, initiating passage 2. For subsequent passages, the MIC of each evolving lineage was determined as described above, and sup^Sg^ concentration used for selection was adjusted to maintain the new MIC, which reached 1% at passage 4. The ALE process was halted after passage 5, when the evolved populations had become resistant to >10% sup^Sg^. Isolation of single colonies from all lineages and passages was performed as described above, and 48 colonies each from passages 3, 4, and 5 were screened for sensitivity to 1% wild-type sup^Sg^.

### Whole genome sequencing and sequence analysis

No reference genome sequence was publically available for *S. antibioticus* NRRL B-1701 on NCBI, an overnight culture of the strain was centrifuged to obtain a 30-50 mg cell pellet. This washed with 1 mL PBS, suspended in 0.5 mL Zymo 1X DNA/RNA Shield, and submitted to Plasmidsaurus for genome sequencing and annotation. To obtain sequences for identifying mutations in Umb-evolved strains, overnight cultures of evolved clones and of the relevant ancestral strains were grown and used for total gDNA extraction with the DNeasy Blood & Tissue kit (Qiagen). Sequencing libraries were constructed using the Nextera DNA Flex Library Prep Kit (Illumina) and sequencing was performed with an Illumina MiniSeq or an Illumina iSeq (300 cycles paired end program on each instrument). Reads were aligned to the *Streptomyces griseus subsp. griseus* NBRC 13350 (genome accession GCA_000010605.1) or *S. antibioticus* NRRL B-1701 reference genome using Geneious. Reads were trimmed using the BBDuk plugin. SNPs were called using Geneious with a minimum coverage cutoff of 5 and a minimum variant frequency of 0.9. SNPs present in the ancestor strain were filtered out of the results for the evolved isolates. Only SNPs present in coding sequences were considered. Sequencing data for evolved isolates and the genome sequence of *S. antibioticus* NRRL B-1701 are available under BioProject PRJNA125743.

### Wall teichoic and teichuronic acid isolation and structural characterization

#### Isolation of wall teichoic acids

Isolation of wall teichoic acids from *Streptomyces* cells was performed essentially as described previously^16^. In a typical preparation, 4 g of cells in a 50 mL conical centrifuge tube were suspended in 10 mL of cold 2 M NaCl by pipetting, 25 g 0.1 mm zirconia/glass beads (BioSpec Products) were added and the tube was cooled to 4 °C. The sample was vortexed in the cold room four times for 2 min at highest speed (Vortex Genie 2), with breaks between the cycles to allow for cooling of the material. Samples were then centrifuged for 1 min at 1000×*g*, the cell debris suspension was removed, and the beads were washed twice using 10 mL 2 M NaCl, with centrifugation. The combined cell suspension was centrifuged for 15 min at 14,000×*g*, the supernatant was discarded, the pellet resuspended in 20 mL of phosphate-buffered saline (137 mM NaCl, 27 mM KCl, 10 mM Na_2_HPO_4_, 1.8 mM KH_2_PO_4_) and centrifuged again. The pellet was washed in a similar manner by resuspension (20 mL each) and centrifugation three times with 2% sodium dodecylsulfate and three times with ultrapure water. The remaining material, resuspended in 20 mL H_2_O, was incubated at 60 °C with shaking, pelleted and resuspended in 20 mL of 15 mM Tris, pH 7.0 buffer with 4 mg trypsin. Following an overnight incubation at 37 °C with shaking, the sample was centrifuged and the pellet washed with (20 mL each) 1 M Tris, pH 7.0, 1 M Tris with 1 M NaCl and 1 M Tris, followed by three washes with ultrapure water. The resulting pellet was suspended in 10 mL cold 10% trichloroacetic acid (TCA) and rocked overnight at 4 °C. The sample was then centrifuged 45 min at 10,000×*g* at 10 °C, the supernatant removed, combined with 1/10 volume of 3 M sodium acetate, pH 5.1, and chilled at 4 °C. To the solution, three volumes of cold 95% ethanol was added and the mixture was incubated at −18 °C for at least 16 h. The sample was then centrifuged 40 min at 10,000×*g* at room temperature, the supernatant carefully removed and the pellet of released wall teichoic acid washed from the tube wall with 2 mL 95% ethanol twice. The suspension was centrifuged 5 min at 16,000×*g* and the pellet washed by centrifugation five times with 95% ethanol (1.5 mL each). The residual ethanol was dried on a centrifugal vacuum evaporator, the pellet dissolved in 0.6 mL ultrapure water and centrifuged 5 min at 16,000×*g* to remove water-insoluble material. The solution was lyophilized and the wall teichoic acids were analyzed as described below.

#### Dephosphorylation

Lyophilized WT *S. griseus* TUA–WTA (1.4 mg) was dissolved in 300 μL cold 48% hydrofluoric acid (HF) on ice and the solution was kept at 4 °C for 30 h. HF was evaporated with a stream of air to ∼30 μL, diluted with ultrapure water to 300 μL and evaporated again. Following dilution to 300 μL, the sample was lyophilized and fractionated using size exclusion chromatography (SEC).

#### Size exclusion chromatography (SEC)

SEC was performed on a Superdex 75 10/300 GL column (Cytiva) connected to an Agilent 1260 Infinity II high-performance liquid chromatography system equipped with a refractive index detector (RID) for monitoring eluting species. The separations of the intact (50 μg) and dephosphorylated (1.4 mg) TUA–WTA material isolated from WT *S. griseus* (dissolved in water) were performed at a flow rate of 0.5 mL/min of 50 mM ammonium acetate buffer (pH 6). A semi-preparative SEC of the WT *S. griseus* TUA/WTA dissolved in the running buffer (two runs of 1.3 mg each) was performed in 100 mM ammonium acetate (pH 6). Because of the shift between RID and fraction collector, each fraction was analyzed by ^1^H NMR and assessed for the presence of signals characteristic for TUA and WTA. Representative early and late fractions were also analyzed by ^1^H,^13^C-HSQC and DOSY. Based on the results, fractions within 10-12 ml elution volumes were pooled as those containing TUA+WTA, and the WTA-only containing fractions from elution volumes 13-15 ml were pooled separately. The structures present in each pool were determined by 1D and 2D NMR, as described below.

#### Mass spectrometry analysis

Mass spectrometry (MS) analysis of the dephosphorylated *S. griseus* TUA material was performed by the matrix-assisted laser desorption/ionization time-of-flight (MALDI-TOF) technique on a Bruker rapifleX mass spectrometer. Mass spectra of the TUA mixed with SDHB matrix (9:1 2,5-dihydroxybenzoic acid:2-hydroxy-5-methoxybenzoic acid) were acquired in the reflector negative mode. The spectrometer was calibrated using angiotensin-I peptide and the spectra were analyzed in Bruker Daltonik FlexControl 3.0 software.

#### NMR spectroscopy

Each NMR sample was prepared from 1–2 mg of dry material that was first dissolved in 200 μL D_2_O (99.9% D, Sigma) with 60 nmol DSS-*d*_6_ (Cambridge Isotope Laboratories) and lyophilized. Following the deuterium exchange, the sample was dissolved in 42 μL D_2_O (99.96% D, Thermo Scientific) and transferred into a 1.7 mm NMR tube. For *S. griseus,* a sample of the SEC-separated higher mass material was also prepared in 97:3 H_2_O:D_2_O solvent to observe signals due to amide hydrogens. NMR data were acquired on a Bruker Avance NEO NMR spectrometer (^1^H, 799.71 MHz) equipped with a 1.7 mm TCI cryoprobe. The pulse programs included in the spectrometer library were used to acquire 1D ^1^H and 2D COSY, TOCSY, NOESY, ROESY, DOSY, ^1^H,^13^C-HSQC, ^1^H,^13^C-HMBC and ^1^H,^13^C-HSQC-TOCSY data. A ^13^C-coupled IPAP-HSQC experiment was acquired to determine ^1^H–^13^C ^1^*J* scalar couplings and confirm the anomeric α/β configurations. Quantitative ^1^H NMR data were acquired with a total recovery delay of 60 s. Mixing time was set to 80 ms for NOESY and 200 ms for ROESY experiments, to 70 and 140 ms for TOCSY experiments, and to 70 ms for HSQC-TOCSY experiment. The DOSY experiments were run using a stimulated echo with longitudinal eddy current delay and bipolar gradient pulses. Gradient strength was varied from 3 to 97% maximum power in 32 linear increments. The diffusion gradient length was 2 ms and diffusion delay (400– 600 ms) was optimized for each sample. The NMR data for *S. griseus* samples were acquired at 313 K, except for the DOSY spectra acquired at 298 K. The NMR data for *S. antibioticus* samples were acquired at 298 K. ^1^H and ^13^C chemical shifts were referenced to the respective DSS signals at 0.00 ppm. NMR data were processed in Topspin 4.0.5, analyzed in AnalysisAssign 3.2.0^43^ and rendered for figures in MestReNova (14.2.3). DOSY reconstruction was done in MestReNova using the peak fit algorithm.

#### Structural analysis of the carbohydrate portion of S. griseus TUA-WTA

NMR spectra of TCA-released material from the WT *S. griseus* cell walls indicated that it contained six distinct glycosyl residues that were labeled A–G (Table S3, Figure S2A). The highest-abundance residues C and E were identified as hexopyranoses bearing acetamido groups. Residue C was found to be a 3-substituted α-GalNAc based on the characteristic chemical shifts and the interruption of ^1^H–^1^H spin system at H-4. Residue E was a 2,3-diacetamido-2,3-dideoxyhexuronic acid in *manno* configuration, given the break of the ^1^H–^1^H spin system at H-2 and an HMBC correlation from H-5 to a carbonyl resonance at 176.6 ppm that shifted to lower field at higher pH. The β anomeric configuration of residue E was confirmed by the anomeric ^1^*J*_HC_ of 163 Hz and ROESY interactions H-1–H-3 and H-1–H-5. A disaccharide repeating unit consisting of residues C and E was established based on the glycosidic linkage signals in the NOESY/ROESY and HMBC spectra (Figure 2A) to be →3)GalNAc-α(1→4)-ManNAc3NAcA-β(1→. Chemical shifts of this disaccharide motif agreed closely with those previously reported for TUAs isolated from *S. lavendulocolor* and *S. griseus*^17,44^. From the chemical shift agreement we inferred that both residues were in the D absolute configuration as reported.

In addition to the major residues C and E, weaker signals were present that belonged to four residues constituting the terminal and linker groups. We found that two non-reducing end residues were present, suggesting irregular termination of the chain. Majority of the polymer (∼70%) was terminated by 4-*O*-acetylated β-ManNAc3NAcA (residue D), while ∼30% of the chains were terminated by α-GalNAc (residue B). According to the HMBC and ROESY correlations (Figure S2A), residue D formed a β(1→3) glycosidic bond with a C-like residue, while residue B was α(1→4) linked to an E-like residue, consistent with an alternate termination of the chain.

On the reducing end of the chain, two distinct glycosyl residues were present, a 3-substituted *G*al*p*NAc (residue A) that was α(1→4) linked to a Glc*p*NAc3NAcA (residue G). Residue G was identified as a 2,3-diacetamido-2,3-dideoxyhexuronic acid in *gluco* configuration owing to an uninterrupted ^1^H–^1^H spin system from H-1 to H-5 and an HMBC correlation from H-5 to a carbonyl resonance at 175.2 ppm that was pH-dependent. The β anomeric configuration of residue G was confirmed by the anomeric ^1^*J*_HC_ of 164 Hz, ^3^*J*_HH_ between H-1 and H-2 of 7.8 Hz, and characteristic ROESY interactions. ^13^C chemical shifts of this A–G disaccharide matched well those reported for a TUA with a →3)-D-GalNAc-α(1→4)-’-GlcNAc3NAcA-β(1→ repeat that was found in *Streptomyces* sp VKM Ac-2537^17^; the matching chemical shifts indicate the d absolute configuration of both residues A and G. Based on a ROESY correlation (Figure S2A), the 3-OH of residue A was substituted by an E-like residue. The terminal residue G was then glycosidically linked by a β(1→4) bond to a ribitol-5-phosphate residue (Rg), as apparent from the ROESY and HMBC spectra (Figure S2A).

NMR analysis of the larger-size TUA-WTA molecule obtained by SEC as well as the dephosphorylated TUA both corroborated the structure of the TUA-like glycan and its linkage to a ribitol-5*P* residue (Figure S2B-D, Table S3). In addition to the major glycosyl residues, a small amount of terminal ManNAc3NAcA was identified (residue K) that was likely the product of limited de-O-acetylation of residue D. A partially resolved residue (Cd) was also identified in the SEC-separated TUA-WTA species that was the aglycon of the non-reducing end residue D. NMR data acquired in 97:3 H_2_O:D_2_O provided spectral information for the amide HN groups (Table S3) and were instrumental in verifying the chemical shift assignments and connectivities for the mono- and diamino-hexoses.

In the dephosphorylated TUA, NMR showed that the reducing-end residue G was β(1→4) linked to a terminal ribitol (residue Rh). MALDI-TOF MS analysis of the dephosphorylated TUA (Figure 2E) was consistent with the TUA structure determined by NMR. Two series of peaks were present in the MALDI spectrum that corresponded to the two polymers with alternative non-reducing ends and with varying numbers of the disaccharide repeating unit (m/z differences of 461.16). Masses corresponding to 2–6 repeats with the 4OAc-ManNAc3NAcA terminus, as well as the de-*O*-acetylated equivalents were present. Similarly, masses corresponding to 1–6 repeats with the GalNAc terminus were detected.

#### Structural analysis of S. griseus WTA

Analysis of polyol phosphates present in the WTA portion of the TCA-released material from the cell walls of WT *S. griseus* was based mainly on the HSQC, HSQC-TOCSY and HMBC spectra that circumvented the overlaps present in the ^1^H–^1^H correlation spectra. The analysis provided individual polyol-*P* units but their connectivities could not be defined using the above experiments. ^1^H,^31^P-HMBC spectrum was also acquired in an attempt to establish sequential connections of the polyols through phosphodiester groups but signal crowding prevented its meaningful analysis.

Based on the HSQC-TOCSY and HMBC correlations as well as characteristic ^1^H/^13^C chemical shifts^21,22,45^ and symmetry considerations, three major polyol-*P* groups were identified in the total TCA hydrolysate as well as the SEC-separated TUA-WTA and WTA material (Table S3): a 1,5-linked Rbo*P* (residue Ra), a 1,3,5-linked Rbo*P* (residue Rb) and a terminal 3-linked Gro*P* (residue Gr). Low-abundance terminal 3-linked Rbo*P* (residue Re) was also found in the lower-mass WTA material (Table S3). The composition of the polyol-*P* residues indicated that *S. griseus* WTA is mainly the poly(Rbo*P*) type and may contain Gro*P* branches or termini. A very low level of lysine modification was also detected in all WTA preparations.

Analysis of the *S. griseus* Δ*utrC* WTA material showed that it predominantly consisted of 1,5-linked Rbo*P* (Ra) with reduced amounts of terminal Gro*P* (Gr) residue (Table S3). The mutant WTA contained a terminal Rbo-5*P* (residue Rc) that was not observed in the WT; however, residue Rc could be analogous to the WT residue Rg that is not substituted with the TUA. The content of branching Rbo*P* (Rb) was very low in the mutant WTA. An additional, unidentified polyol-*P* (residue Rd) was also present.

#### Structural analysis of S. antibioticus TUA-WTA and WTA

Based on the analysis of NMR spectra, the TCA-released material from WT *S. antibioticus* contained seven distinct glycosyl residues, labeled A–G (Table S7, Figure S7). A subset of the residues (C, D, F, G) was each found to be a 2,3-diacetamido-2,3-dideoxyhexuronic acid in *gluco* configuration owing to an uninterrupted ^1^H–^1^H spin system from H-1 to H-5 and an HMBC correlation from H-5 to a carbonyl resonance at ∼173 ppm (low pH).

The most abundant residue C formed a β(1→4) linked homo-oligomer, →4)-Glc*p*NAc3NAcA-β(1→, as was apparent from the HMBC and ROESY inter-residue correlations. Residue F was the non-reducing end of the oligomer bearing a 4-O-acetyl group. On the reducing end, a partially resolved GlcNAc3NAcA (residue D) connected the chain to a terminal Glc*p*NAc3NAcA with distinct chemical shifts (residue G); residue D then linked the oligomer to a Rbo-5P residue (Rg) via a β(1→4) glycosidic bond. The absolute configuration of the GlcNAc3NAcA residues has not been determined. To the best of our knowledge, a homopolymer of GlcNAc3NAcA has not been reported previously.

The remaining three residues (A, B, E) were identified as terminal β-Glc*p* residues glycosidically linked to the polyol-P backbone of the WTA (Figure S7A,D). The most abundant residue A connected via a β(1→4) bond to 1,5-linked ribitol-5P unit (residue Rf). A small proportion of the Glc residues were 3-O-methylated (residue B). A similar glucosylated Rbo-5P polymer has been reported in WTA from other *Streptomyces*^21,44^. The WTA also contained unsubstituted 1,5-linked ribitol-5P residues (Ra). A minor residue C was then β(1→4) glycosidically linked to another polyol-P that has not been identified.

The spectra of *S. antibioticus* Δ*utrC* TCA-released material lacked signals of the TUA-like residues C, D, F and G, and contained only signals of glycosyl residues A, B and E, together with ribitol-P residues Ra and Rf (Table S8, Figure S7A,B). Additionally, similar to what was observed for *S. griseus* Δ*utrC*, a terminal Rbo-5P was present (residue Rc) that could be analogous to the WT residue Rg not substituted with the TUA-like chain in the mutant.

### Bacterial pull-down assay and whole cell proteome analysis

For detection of proteins that bind to Umb target cells, spores of target organisms *S. griseus* and *S. antibioticus* were cultured in 30 mL TSBY medium for 20 h. Cells were pelleted by spinning at 3,000 g for 5 min, washed twice with TSBY medium, and resuspended in TSBY medium to an OD_600_ of approximately 4. A 500 μL cell aliquot was mixed with 500 μL of sup^Sc^ (from *S. coelicolor* Δ*umbC1-3*), sup^Sg^ (from *S. griseus* Δ*umbC*), a 1:1 (v/v) mixture of sup^Sc^ and sup^Sg^, or PBS (negative control) in a 1.5 mL microcentrifuge tube. Samples were incubated at RT with continuous rotation for 1.5 h. Then cells were pelleted by spinning at 3,000 g for 5 min, washed three times with PBS, resuspended in 400 uL lysis buffer (8 M urea, 75 mM NaCl, 50 mM Tris-HCl, pH 8.2) and lysed by sonication. After spinning at 20,000 g for 20 min, 20 μL of resulting supernatant was used to determine protein concentration via the Pierce^TM^ BCA Protein Assay Kit (Thermo Fisher Scientific). Meanwhile, 100 μL of protein solutions were transferred to a new 1.5 mL microcentrifuge tube and incubated with 5 mM Dithiothreitol (DTT) at 56 ℃ for 25 min for protein reduction. After cooling to room temperature, 14 mM iodoacetamide was added for protein alkylation and the samples were incubated in the dark at room temperature for 30 min. The reaction was quenched by adding additional 5 mM DTT and incubating 15 min in the dark at room temperature. The mixture was diluted 5-fold with 50 mM Tris-HCl (pH 8.2) to reduce the urea concentration to 1.6 M and 1 mM calcium chloride was then added. Approximately 20 μg protein (as determined by BCA assay) was mixed with trypsin (Promega) at 4 μg/mL and incubated at 37 ℃ for 16 h. Digestion was halted by adding 0.4% trifluoroacetic acid (TFA) to lower the pH below 2.

Trypsin-digested peptides were purified using in-house constructed stop-and-go-extraction tips (StageTips)^46^ embedded with 6 layers of Empore^TM^ styrenedivinylbenzene-reverse phase sulfonate (SDB-RPS) disks (Sigma). StageTips were conditioned sequentially with 50 μL of 100% methanol, 100% acetonitrile (ACN), 75%ACN and 5% ammonium hydroxide, 75% ACN and 0.5% acetic acid, and 0.1% TFA. Digested peptide samples were loaded onto StageTips and washed sequentially with 50 μL of 0.1% TFA, 75% ACN and 0.5% acetic acid, 0.5% acetic acid, then eluted with 60 μL of 75%ACN and 5% ammonium hydroxide. Peptides were dried using a SpeedVac^TM^ vacuum concentrator (Thermo Fisher Scientific) and resuspended in 5% ACN with 0.1% formic acid and analyzed by LC-MS/MS on a Lumos Fusion Orbitrap mass spectrometer (Thermo Scientific) as previously described^47^.

For analysis of proteins in sup^Sc^ and sup^Sg^ (“reference”), 100 μL of protein solution was mixed with 25 μL 100% trichloroacetic acid (TCA) and incubated on ice for 30 min to precipitate proteins. After centrifugation at 20,000 x g for 5 min, the protein pellet was washed once with 20% TCA and twice with 100% acetone. The dried protein samples were resuspended in lysis buffer (8 M urea, 75 mM NaCl, 50 mM Tris-HCl, pH 8.2) and subsequent reduction, alkylation, trypsin digestion, and peptide purification and mass spectrometry were performed as described above.

Mass spectrometry data were analyzed using MaxQuant^48^. Data were filtered to remove proteins identified in the negative control, where cells were incubated with PBS instead of supernatant. The proportion of mass spectral counts for proteins in “reference” and “cell-associated” groups was then calculated. Two biological replicates were included in each group.

### Purification of UmbA4–UmbB

*S. coelicolor* Δ*umbC1-3* pSET152–*attB*::*umbA4*–2xGGGGS–3xK–6xH was used for purification of UmbA4–UmbB complexes. Spores were cultured in 30 mL 2xYT broth (Research Products International) for 24 h, then back diluted to an OD_600_ of 0.01 in 100 mL 2xYT broth for a total combined culture volume of 4 L. Cultures were incubated for approximately 28 h until OD_600_ reached 5-6. Cells were pelleted by centrifugation at 21,000 g for 30 min, and the supernatant was filtered using Rapid-Flow^TM^ Filter Units (PES Membrane, 0.45 um; Thermo Fisher Scientific). A total of 3.5 L of supernatant was combined with 700 mL of 5xPBS buffer and incubated with approximately 10 mL bed volume TALON cobalt resin (Takara) under continuous stirring at 4 ℃ for 4 h. The resin was separated from supernatant by centrifugation at 300 g for 3 min and packed into two Econo-Column® Chromatography Columns (2.5 × 30 cm; Bio-Rad). The approximately 4 L supernatant was loaded onto the columns, followed by washing with PBS buffer. Bound proteins were eluted with PBS containing 500 mM imidazole. Eluted fractions were concentrated using a 30 kDa cut-off Amicon concentrator to a final volume of 1.1 mL. The protein sample was further purified by AKATA FPLC using a Superose 6 Increase 10/300 GL column (GE healthcare) equilibrated in sizing buffer (250 mM NaCl, 2.7 mM KCl, 8 mM Na_2_HPO_4_, and 2 mM KH_2_PO_4_, pH 7.2). Each fraction was assessed for purity by SDS-PAGE with Coomassie blue staining. Fractions with high purity and concentration were further concentrated and used for Cryo-EM or cell binding assays.

### CryoEM sample preparation, data collection and data processing

For structure determination of UmbA4–UmbB complexes in the absence of bound TUA, three μl of 3 mg/ml purified protein complex with 2 mM 3-[(3-Cholamidopropyl)dimethylammonio]-2-hydroxy-1-propanesulfonate (CHAPSO) were loaded onto freshly glow discharged R 2/2 UltrAuFoil grids, prior to plunge freezing using a vitrobot MarkIV (ThermoFisher Scientific) with a blot force of 0 and 5 sec blot time at 100 % humidity and 22°C. Cryo-EM analysis revealed that each purified fraction contained various levels of UmbA4 alone, UmbA4 complexed with UmbB (UmbB1-3 could not be distinguished), proteolytic products containing the C-terminal lectin domain of Umb4 without N-terminal trypsin-like domain and other unidentified protein particles. To obtain a structure of UmbA4– UmbB, 2,380 micrographs of promising fractions were collected using Leginon^49^ at an accelerating voltage of 200kV with Gatan K3 direct electron detector, nominal magnification 45,000x, a calibrated pixel size of 0.89 Å, exposure time of 5 s and a total fluence of 47.3 e/Å^2^. Movie frame alignment, estimation of the microscope contrast-transfer function parameters, particle picking and extraction were carried out using in cryoSPARC^50^. 2D classification, ab- initio reconstruction, heterogeneous 3D refinement and non-uniformed 3D refinement was performed in cryoSPARC. The final 3D refinement of UmbA4 protein was carried out in cryoSPARC, leading to a map with resolution of 4.3 Å comprising 200,137 particles.

To determine the structure of UmbA4 bound to TUA, 50 uM protein and 1 mM teichuronic acid (TUA), purified via dephosphorylation and SEC from *S. griseus* TUA–WTA, were incubated in buffer containing 10 mM Na_2_HPO_4_, 1.8 mM KH_2_PO_4_ pH 7.4, 250 mM NaCl and 2.7 mM KCl for 60 min at 4 °C. To prepare the CryoEM UmbA4–TUA sample grids, three microliters of 3.5 mg/ml UmbA4–TUA with 2 mM CHAPSO were loaded onto freshly glow discharged R 2/2 UltrAuFoil grids, prior to plunge freezing using a vitrobot MarkIV (ThermoFisher Scientific) with a blot force of 0 and 5 sec blot time at 100 % humidity and 22°C. For structure determination of the UmbA4–TUA complexes, data were acquired using an FEI Titan Krios transmission electron microscope operated at 300 kV and equipped with a Gatan K3 direct detector and Gatan Quantum GIF energy filter, operated in zero-loss mode with a slit width of 20 eV. The dose rate was adjusted to 7 counts/pixel/s, and each movie was acquired in 80 frames of 50 ms. Automated data collection was carried out using Leginon^49^ at a nominal magnification of 105,000x with a pixel size of 0.835 Å. The nominal defocus values were set from -0.2 and -1.6 μm and a total of 9,729 movies were collected. Movie frame alignment, estimation of the microscope contrast-transfer function parameters, initial particle picking and extraction were carried out using Warp^51^. Initially, we employed single-particle image analysis with no imposed symmetry to obtain cryo-EM structures of the UmbA4 filaments interacting with TUA polymers. First, the particles picked using Warp were averaged using 2D classification to generate ab-initio reconstruction in cryoSPARC^50^. Then one round of 3D refinement was carried out in C1 symmetry without helical parameters and the reconstructed maps were used to estimate the initial helical symmetry parameters of UmbA4–TUA complex. The helical parameters for the filament were first approximated empirically in real space using Chimera^52^, and these parameters were then refined using three-dimensional helical refinement with symmetry searches in cryoSPARC. For filament segments picking, micrographs were imported into cryoSPARC. The filament tracer module was used to pick filament segments from micrographs with fibril diameter 60-160 Å and segment overlap of 0.4 x diameter which is equivalent to a helical rise. 5,747,176 filament segments were extracted in 168 pixel (280.6 Å) boxes, covering roughly six unique UmbA4-TUA subunits per box segments. Filament segments were subjected to reference-free 2D classification in cryoSPARC, and six well-defined filament class averages were selected for two rounds of reference-free ab-initio reconstruction and heterogenous 3D refinements without symmetry. 1,492,839 of the best filament segments were used for 3D helical refinement with a rise of 34.2 Å, helical twist of 208.3° and C1 symmetry in cryoSPARC. The selected dataset was transferred from cryoSPARC to Relion format using the pyem program package and particle images were subjected to the Bayesian polishing procedure implemented in Relion^53^ during which particles were re-extracted with a box size of 280 pixels and a pixel size of 1.0 Å. To eliminate poor quality particles, another round of reference-free 2D classification and ab-initio reconstruction in cryoSPARC was used to re-classify the data and the generated models were used as references for heterogeneous 3D refinement. Helical symmetry search was then carried out again in cryoSPARC. The final 3D refinements of the TUA molecules bound UmbA4 complex structure were carried out using 3D helical refinement with a rise of 33.13 Å, helical twist of 208.86° and per-particle defocus refinement in cryoSPARC to yield the reconstruction at 3.3 Å resolution comprising 1,182,274 particles. Local resolution estimation, filtering, and sharpening were carried out using cryoSPARC. Reported resolutions are based on the gold-standard Fourier shell correlation (FSC) of 0.143 criterion and Fourier shell correlation curves were corrected for the effects of soft masking by high-resolution noise substitution ^54,55^. The details of the image processing procedure are summarized in Figure S3E,K.

### Model building and refinement

The initial model of UmbA4 and UmbB was generated from the AlphaFold3^56^. UCSF Chimera^52^ and Coot^57^ were used to fit atomic models into the cryoEM maps. Carbohydrate models were built and refined into density using Rosetta. Building off previous carbohydrate modelling tools [https://pubmed.ncbi.nlm.nih.gov/30344107/], Rosetta parameter files for each of the five unique sugars were manually built in Rosetta. An initial model was constructed using knowledge of the placement of the reducing end: the carbohydrate chain was grown by adding two sugars at a time and refining the growing model into density. With the initial carbohydrate model built, we then further refined the model of the complete assembly. Using the helical symmetry present in the cryoEM map, we symmetrically refined both protein and carbohydrate together in the model using Rosetta’s *relax*, yielding the final complex. The final model contains UmbA4 residues 33-705 and TUA glycans 1-12. The detail refinement statistics are summarized in Table S4.

### Bioinformatics analysis

#### Gene neighborhood analysis of the WTA loci

We compiled a protein database comprising 20 selected *Streptomyces* genomes, along with several non-*Streptomyces* genomes from other Actinobacteria as outgroups. We initiated multiple PSI-BLAST searches^58^ using several proteins in the Utr locus, and nearby WTA biosynthetic proteins including SGR_4957 (TagF), SGR_4943 (UtrC), SGR_4956 (TarIJ), and two adjacently encoded ribosomal proteins to use as locus placement anchors, SGR_4953 (Ribosomal protein L21p), and SGR_4954 (Ribosomal protein L27). These searches employed a stringent cutoff e-value of 0.005. For each identified homolog, we extracted and analyzed the surrounding genomic context, encompassing 15 upstream and 15 downstream neighboring genes. We found that they typically occupied the same genomic locus, and these were unique to each genome. Proteins recovered from these loci were systematically grouped into major clusters based on sequence similarity using the BLASTCLUST program (https://ftp.ncbi.nih.gov/blast/documents/blastclust.html). Subsequently, we annotated these protein clusters by identifying their constituent domains through hmmscan searches^59^ against Pfam^60^ and our curated domain profiles. Additionally, to predict the function of domain families that are not annotated, we generated the structural models using AlphaFold^56^ and identified the distant homologs using the DALI program^61^.

#### Identification and domain analysis of Streptomyces UmbA proteins

We initially compiled a comprehensive dataset consisting of 11,223 annotated *Streptomycetaceae* genomes retrieved from the NCBI GenBank database^62^, of which 10,618 belonged specifically to the genus *Streptomyces*. Using the inactive trypsin domain of UmbA (residues 21–230 of CAC36720.1) as a query, PSI-BLAST searches^58^ were conducted against this custom genome database with a stringent E-value cutoff of 0.005. This search yielded 9,455 sequences harboring the inactive trypsin domain (UmbA homologs).

Domain architecture annotations were subsequently performed using the hmmscan program^59^, querying both the Pfam profile database^60^ and a custom-curated domain profile database. This analysis highlighted a significant association between the inactive trypsin domain and various membrane-localization domains positioned in the C-terminal regions, including diverse β-propeller domains, ricinB-lectin, and bulb-lectin domains. To further analyze these membrane-localization regions, we extracted the corresponding C-terminal sequences larger than 20 amino acids, resulting in a dataset of 9192 sequences (Dataset 1). Additionally, we observed instances in which the inactive trypsin domain (UmbA) and the associated lectin domains reside in separate but neighboring genes. To systematically retrieve these occurrences, neighboring genes (five upstream and five downstream) relative to each UmbA-containing gene were extracted. These neighboring sequences were subjected to similarity searches against the previously extracted C-terminal UmbA regions using BLASTP. Sequences displaying significant similarity to any of these regions were retained, resulting in an additional set of 667 sequences (Dataset 2). Combining these two datasets resulted in a consolidated dataset comprising 9,859 unique sequences.

To achieve a comprehensive identification and classification of domain families within these 9,859 sequences, we employed the following strategy:

1. Network clustering analysis was conducted using the CLANS program^63^, which employs the Fruchterman and Reingold force-directed layout algorithm based on significant high-scoring segment pairs (HSPs) identified through all-against-all BLASTP searches (P-value: 0.0001). The sequences, represented as nodes, and the edges between are attractive forces which are proportional to the negative logarithm of the alignment significance values. This approach facilitated the identification of major sequence clusters among these 9,859 sequences.
2. For each densely connected subgraph (cluster), multiple sequence alignment was generated using either KALIGN^64^ or MUSCLE^65^. Representative sequences from each alignment were selected for structural modeling using AlphaFold^56,66^. Domain boundaries were delineated through analyses of inter-residue distance matrices generated from predicted structural models.
3. The identified domain sequences were further clustered using the CLANS program to precisely define distinct domain families, yielding a total of 41 major families.
4. These domain families were subsequently grouped into major fold superfamilies through sequence profile comparisons using HHsearch^67^ and structural similarity comparisons using the DALI server^61^.

Through this comprehensive analysis, we identified 41 distinct domain families that belong to 12 overarching fold superfamilies. These include 12 families of five-bladed β-propellers (5BP1 to 5BP12), four families of six-bladed β-propellers (6BP1 to 6BP4), seven families of seven-bladed β-propellers (7BP1 to 7BP7), nine ricinB-lectin families (RB1 to RB9), one bulb-lectin family, one jellyroll-like fold family, two GDSL hydrolase domain families, one laminin_G_3 family, one GDPD domain family, one CBM_4_9 family, one newly identified immunoglobulin-like domain family (BPA-Ig), and a novel β-sandwich fold family designated TAC4.

### Statistics and reproducibility

Significance of differences in growth yields from supernatant toxicity assays and in protein abundances from whole cell proteome analyses were determined using two-tailed *t*-tests. Tests were performed using Graphpad Prism. For bacterial growth assays, bacterial pull-down assays, and cell binding assays, the number of replicates collected from independent cultures grown in parallel on a single day are indicated in corresponding methods. Each experiment was also replicated at least once on separate days with at least two additional cultures. Statistical methods were not used to predetermine sample size, and randomization and blinding were not employed.

## Notes

### Competing Interest Statement

The authors have declared no competing interest.

## References

1. Stone, E., Campbell, K., Grant, I., and McAuliffe, O. (2019). Understanding and Exploiting Phage-Host Interactions. Viruses 11. 10.3390/v11060567.

2. van Dalen, R., Peschel, A., and van Sorge, N.M. (2020). Wall Teichoic Acid in Staphylococcus aureus Host Interaction. Trends Microbiol 28, 985–998. 10.1016/j.tim.2020.05.017.

3. Wu, X., Han, J., Gong, G., Koffas, M.A.G., and Zha, J. (2021). Wall teichoic acids: physiology and applications. FEMS Microbiol Rev 45. 10.1093/femsre/fuaa064.

4. Cascales, E., Buchanan, S.K., Duche, D., Kleanthous, C., Lloubes, R., Postle, K., Riley, M., Slatin, S., and Cavard, D. (2007). Colicin biology. Microbiol Mol Biol Rev 71, 158–229. 71/1/158 [pii] 10.1128/MMBR.00036-06.

5. Alam, K., Mazumder, A., Sikdar, S., Zhao, Y.M., Hao, J., Song, C., Wang, Y., Sarkar, R., Islam, S., Zhang, Y., and Li, A. (2022). Streptomyces: The biofactory of secondary metabolites. Front Microbiol 13, 968053. 10.3389/fmicb.2022.968053.

6. Barka, E.A., Vatsa, P., Sanchez, L., Gaveau-Vaillant, N., Jacquard, C., Meier-Kolthoff, J.P., Klenk, H.P., Clement, C., Ouhdouch, Y., and van Wezel, G.P. (2016). Taxonomy, Physiology, and Natural Products of Actinobacteria. Microbiol Mol Biol Rev 80, 1–43. 10.1128/MMBR.00019-15.

7. Hopwood, D.A. (2007). Streptomyces in Nature and Medicine (Oxford University Press).

8. McCormick, J.R., and Flardh, K. (2012). Signals and regulators that govern Streptomyces development. FEMS Microbiol Rev 36, 206–231. 10.1111/j.1574-6976.2011.00317.x.

9. Kinkel, L.L., Schlatter, D.C., Xiao, K., and Baines, A.D. (2014). Sympatric inhibition and niche differentiation suggest alternative coevolutionary trajectories among Streptomycetes. ISME J 8, 249–256. 10.1038/ismej.2013.175.

10. Zhao, Q., Bertolli, S., Park, Y.J., Tan, Y., Cutler, K.J., Srinivas, P., Asfahl, K.L., Fonesca-Garcia, C., Gallagher, L.A., Li, Y., et al. (2024). Streptomyces umbrella toxin particles block hyphal growth of competing species. Nature 629, 165–173. 10.1038/s41586-024-07298-z.

11. Ruhe, Z.C., Low, D.A., and Hayes, C.S. (2020). Polymorphic Toxins and Their Immunity Proteins: Diversity, Evolution, and Mechanisms of Delivery. Annu Rev Microbiol 74, 497–520. 10.1146/annurev-micro-020518-115638.

12. Riley, M.A. (1998). Molecular mechanisms of bacteriocin evolution. Annu Rev Genet 32, 255–278. 10.1146/annurev.genet.32.1.255.

13. Brown, S., Xia, G., Luhachack, L.G., Campbell, J., Meredith, T.C., Chen, C., Winstel, V., Gekeler, C., Irazoqui, J.E., Peschel, A., and Walker, S. (2012). Methicillin resistance in Staphylococcus aureus requires glycosylated wall teichoic acids. Proc Natl Acad Sci U S A 109, 18909–18914. 10.1073/pnas.1209126109.

14. Sobhanifar, S., Worrall, L.J., King, D.T., Wasney, G.A., Baumann, L., Gale, R.T., Nosella, M., Brown, E.D., Withers, S.G., and Strynadka, N.C. (2016). Structure and Mechanism of Staphylococcus aureus TarS, the Wall Teichoic Acid beta-glycosyltransferase Involved in Methicillin Resistance. PLoS pathogens 12, e1006067. 10.1371/journal.ppat.1006067.

15. Xia, G., Maier, L., Sanchez-Carballo, P., Li, M., Otto, M., Holst, O., and Peschel, A. (2010). Glycosylation of wall teichoic acid in Staphylococcus aureus by TarM. J Biol Chem 285, 13405–13415. 10.1074/jbc.M109.096172.

16. Kho, K., and Meredith, T.C. (2018). Extraction and Analysis of Bacterial Teichoic Acids. Bio Protoc 8, e3078. 10.21769/BioProtoc.3078.

17. Tul’skaya, E.M., Shashkov, A.S., Streshinskaya, G.M., Senchenkova, S.N., Potekhina, N.V., Kozlova, Y.I., and Evtushenko, L.I. (2011). Teichuronic and teichulosonic acids of actinomycetes. Biochemistry (Mosc) 76, 736–744. 10.1134/S0006297911070030.

18. Ward, J.B. (1981). Teichoic and teichuronic acids: biosynthesis, assembly, and location. Microbiol Rev 45, 211–243. 10.1128/mr.45.2.211-243.1981.

19. Gassner, G.T., Dickie, J.P., Hamerski, D.A., Magnuson, J.K., and Anderson, J.S. (1990). Teichuronic acid reducing terminal N-acetylglucosamine residue linked by phosphodiester to peptidoglycan of Micrococcus luteus. J Bacteriol 172, 2273–2279. 10.1128/jb.172.5.2273-2279.1990.

20. Ward, J.B., and Curtis, C.A. (1982). The biosynthesis and linkage of teichuronic acid to peptidoglycan in Bacillus licheniformis. Eur J Biochem 122, 125–132. 10.1111/j.1432-1033.1982.tb05857.x.

21. Naumova, I.B., and Shashkov, A.S. (1997). Anionic polymers in cell walls of gram-positive bacteria. Biochemistry (Mosc) 62, 809–840.

22. Uchikawa, K., Sekikawa, I., and Azuma, I. (1986). Structural studies on teichoic acids in cell walls of several serotypes of Listeria monocytogenes. Journal of biochemistry 99, 315–327. 10.1093/oxfordjournals.jbchem.a135486.

23. Soldo, B., Lazarevic, V., and Karamata, D. (2002). tagO is involved in the synthesis of all anionic cell-wall polymers in Bacillus subtilis 168. Microbiology (Reading) 148, 2079–2087. 10.1099/00221287-148-7-2079.

24. Soldo, B., Lazarevic, V., Pagni, M., and Karamata, D. (1999). Teichuronic acid operon of Bacillus subtilis 168. Molecular microbiology 31, 795–805. 10.1046/j.1365-2958.1999.01218.x.

25. Bonnardel, F., Kumar, A., Wimmerova, M., Lahmann, M., Perez, S., Varrot, A., Lisacek, F., and Imberty, A. (2019). Architecture and Evolution of Blade Assembly in beta-propeller Lectins. Structure 27, 764–775 e763. 10.1016/j.str.2019.02.002.

26. Jones, G.H., Paget, M.S., Chamberlin, L., and Buttner, M.J. (1997). Sigma-E is required for the production of the antibiotic actinomycin in Streptomyces antibioticus. Molecular microbiology 23, 169–178. 10.1046/j.1365-2958.1997.2001566.x.

27. Mispelaere, M., De Rop, A.S., Hermans, C., De Maeseneire, S.L., Soetaert, W.K., De Mol, M.L., and Hulpiau, P. (2024). Whole genome-based comparative analysis of the genus Streptomyces reveals many misclassifications. Appl Microbiol Biotechnol 108, 453. 10.1007/s00253-024-13290-4.

28. Westman, E.L., McNally, D.J., Charchoglyan, A., Brewer, D., Field, R.A., and Lam, J.S. (2009). Characterization of WbpB, WbpE, and WbpD and reconstitution of a pathway for the biosynthesis of UDP-2,3-diacetamido-2,3-dideoxy-D-mannuronic acid in Pseudomonas aeruginosa. J Biol Chem 284, 11854–11862. 10.1074/jbc.M808583200.

29. Pereira, M.P., and Brown, E.D. (2004). Bifunctional catalysis by CDP-ribitol synthase: convergent recruitment of reductase and cytidylyltransferase activities in Haemophilus influenzae and Staphylococcus aureus. Biochemistry 43, 11802–11812. 10.1021/bi048866v.

30. Willett, J.L., Gucinski, G.C., Fatherree, J.P., Low, D.A., and Hayes, C.S. (2015). Contact-dependent growth inhibition toxins exploit multiple independent cell-entry pathways. Proc Natl Acad Sci U S A 112, 11341–11346. 10.1073/pnas.1512124112.

31. Willett, J.L., Ruhe, Z.C., Goulding, C.W., Low, D.A., and Hayes, C.S. (2015). Contact-Dependent Growth Inhibition (CDI) and CdiB/CdiA Two-Partner Secretion Proteins. J Mol Biol 427, 3754–3765. 10.1016/j.jmb.2015.09.010.

32. Houser, J., Komarek, J., Cioci, G., Varrot, A., Imberty, A., and Wimmerova, M. (2015). Structural insights into Aspergillus fumigatus lectin specificity: AFL binding sites are functionally non-equivalent. Acta Crystallogr D Biol Crystallogr 71, 442–453. 10.1107/S1399004714026595.

33. Liu, M., Cheng, X., Wang, J., Tian, D., Tang, K., Xu, T., Zhang, M., Wang, Y., and Wang, M. (2020). Structural insights into the fungi-nematodes interaction mediated by fucose-specific lectin AofleA from Arthrobotrys oligospora. Int J Biol Macromol 164, 783–793. 10.1016/j.ijbiomac.2020.07.173.

34. Lahooti, M., and Harwood, C.R. (1999). Transcriptional analysis of the Bacillus subtilis teichuronic acid operon. Microbiology (Reading) 145 ( *Pt 12*), 3409–3417. 10.1099/00221287-145-12-3409.

35. Meireles, D., Pombinho, R., Carvalho, F., Sousa, S., and Cabanes, D. (2020). Listeria monocytogenes Wall Teichoic Acid Glycosylation Promotes Surface Anchoring of Virulence Factors, Resistance to Antimicrobial Peptides, and Decreased Susceptibility to Antibiotics. Pathogens 9. 10.3390/pathogens9040290.

36. Weidenmaier, C., and Peschel, A. (2008). Teichoic acids and related cell-wall glycopolymers in Gram-positive physiology and host interactions. Nat Rev Microbiol 6, 276-287. 10.1038/nrmicro1861.

37. Winstel, V., Xia, G., and Peschel, A. (2014). Pathways and roles of wall teichoic acid glycosylation in Staphylococcus aureus. Int J Med Microbiol 304, 215–221. 10.1016/j.ijmm.2013.10.009.

38. Xu, Z., Zhang, H., Sun, X., Liu, Y., Yan, W., Xun, W., Shen, Q., and Zhang, R. (2019). Bacillus velezensis Wall Teichoic Acids Are Required for Biofilm Formation and Root Colonization. Applied and environmental microbiology 85. 10.1128/AEM.02116-18.

39. Bektas, S., and Kaptan, E. (2024). Microbial lectins as a potential therapeutics for the prevention of certain human diseases. Life Sci 346, 122643. 10.1016/j.lfs.2024.122643.

40. Zhao, Q., Xie, H., Peng, Y., Wang, X., and Bai, L. (2017). Improving acarbose production and eliminating the by-product component C with an efficient genetic manipulation system of Actinoplanes sp. SE50/110. Synth Syst Biotechnol 2, 302–309. 10.1016/j.synbio.2017.11.005.

41. Dubeau, M.P., Ghinet, M.G., Jacques, P.E., Clermont, N., Beaulieu, C., and Brzezinski, R. (2009). Cytosine deaminase as a negative selection marker for gene disruption and replacement in the genus Streptomyces and other actinobacteria. Applied and environmental microbiology 75, 1211–1214. 10.1128/AEM.02139-08.

42. Bierman, M., Logan, R., O’Brien, K., Seno, E.T., Rao, R.N., and Schoner, B.E. (1992). Plasmid cloning vectors for the conjugal transfer of DNA from Escherichia coli to Streptomyces spp. Gene 116, 43–49. 10.1016/0378-1119(92)90627-2.

43. Skinner, S.P., Fogh, R.H., Boucher, W., Ragan, T.J., Mureddu, L.G., and Vuister, G.W. (2016). CcpNmr AnalysisAssign: a flexible platform for integrated NMR analysis. J Biomol NMR 66, 111–124. 10.1007/s10858-016-0060-y.

44. Shashkov, A.S., Kozlova, Y.I., Streshinskaya, G.M., Kosmachevskaya, L.N., Bueva, O.V., Evtushenko, L.I., and Naumova, I.B. (2001). The Carbohydrate-containing Cell-Wall Polymers of Certain Species from the Cluster “Streptomyces lavendulae”. Microbiology (Moscow) 70, 413–421.

45. Tarellia, E., and Coley, J. (1979). 13C-n.m.r. spectra of some ribitol teichoic acids. Carbohydrate Research 75, 31–37.

46. Rappsilber, J., Mann, M., and Ishihama, Y. (2007). Protocol for micro-purification, enrichment, pre-fractionation and storage of peptides for proteomics using StageTips. Nature protocols 2, 1896–1906. 10.1038/nprot.2007.261.

47. Ting, S.Y., LaCourse, K.D., Ledvina, H.E., Zhang, R., Radey, M.C., Kulasekara, H.D., Somavanshi, R., Bertolli, S.K., Gallagher, L.A., Kim, J., et al. (2022). Discovery of coordinately regulated pathways that provide innate protection against interbacterial antagonism. Elife 11. 10.7554/eLife.74658.

48. Tyanova, S., Temu, T., and Cox, J. (2016). The MaxQuant computational platform for mass spectrometry-based shotgun proteomics. Nature protocols 11, 2301–2319. 10.1038/nprot.2016.136.

49. Suloway, C., Pulokas, J., Fellmann, D., Cheng, A., Guerra, F., Quispe, J., Stagg, S., Potter, C.S., and Carragher, B. (2005). Automated molecular microscopy: the new Leginon system. J Struct Biol 151, 41–60. 10.1016/j.jsb.2005.03.010.

50. Punjani, A., Rubinstein, J.L., Fleet, D.J., and Brubaker, M.A. (2017). cryoSPARC: algorithms for rapid unsupervised cryo-EM structure determination. Nature methods 14, 290–296. 10.1038/nmeth.4169.

51. Tegunov, D., and Cramer, P. (2019). Real-time cryo-electron microscopy data preprocessing with Warp. Nature methods 16, 1146–1152. 10.1038/s41592-019-0580-y.

52. Pettersen, E.F., Goddard, T.D., Huang, C.C., Couch, G.S., Greenblatt, D.M., Meng, E.C., and Ferrin, T.E. (2004). UCSF Chimera--a visualization system for exploratory research and analysis. J Comput Chem 25, 1605–1612. 10.1002/jcc.20084.

53. Zivanov, J., Nakane, T., and Scheres, S.H.W. (2019). A Bayesian approach to beam-induced motion correction in cryo-EM single-particle analysis. IUCrJ 6, 5–17. 10.1107/S205225251801463X.

54. Chen, S., McMullan, G., Faruqi, A.R., Murshudov, G.N., Short, J.M., Scheres, S.H., and Henderson, R. (2013). High-resolution noise substitution to measure overfitting and validate resolution in 3D structure determination by single particle electron cryomicroscopy. Ultramicroscopy 135, 24–35. 10.1016/j.ultramic.2013.06.004.

55. Rosenthal, P.B., and Henderson, R. (2003). Optimal determination of particle orientation, absolute hand, and contrast loss in single-particle electron cryomicroscopy. J Mol Biol 333, 721–745. 10.1016/j.jmb.2003.07.013.

56. Abramson, J., Adler, J., Dunger, J., Evans, R., Green, T., Pritzel, A., Ronneberger, O., Willmore, L., Ballard, A.J., Bambrick, J., et al. (2024). Accurate structure prediction of biomolecular interactions with AlphaFold 3. Nature 630, 493–500. 10.1038/s41586-024-07487-w.

57. Emsley, P., Lohkamp, B., Scott, W.G., and Cowtan, K. (2010). Features and development of Coot. Acta Crystallogr D Biol Crystallogr 66, 486–501. 10.1107/S0907444910007493.

58. Altschul, S.F., Madden, T.L., Schaffer, A.A., Zhang, J., Zhang, Z., Miller, W., and Lipman, D.J. (1997). Gapped BLAST and PSI-BLAST: a new generation of protein database search programs. Nucleic Acids Res 25, 3389–3402.

59. Eddy, S.R. (2011). Accelerated Profile HMM Searches. PLoS computational biology 7, e1002195. 10.1371/journal.pcbi.1002195.

60. Mistry, J., Chuguransky, S., Williams, L., Qureshi, M., Salazar, G.A., Sonnhammer, E.L.L., Tosatto, S.C.E., Paladin, L., Raj, S., Richardson, L.J., et al. (2021). Pfam: The protein families database in 2021. Nucleic Acids Res 49, D412–D419. 10.1093/nar/gkaa913.

61. Holm, L. (2020). Using Dali for Protein Structure Comparison. Methods in molecular biology (Clifton, N.J 2112, 29–42. 10.1007/978-1-0716-0270-6_3.

62. Sayers, E.W., Cavanaugh, M., Clark, K., Pruitt, K.D., Schoch, C.L., Sherry, S.T., and Karsch-Mizrachi, I. (2022). GenBank. Nucleic Acids Res 50, D161–D164. 10.1093/nar/gkab1135.

63. Frickey, T., and Lupas, A. (2004). CLANS: a Java application for visualizing protein families based on pairwise similarity. Bioinformatics 20, 3702–3704. 10.1093/bioinformatics/bth444.

64. Lassmann, T. (2019). Kalign 3: multiple sequence alignment of large data sets. Bioinformatics 36, 1928–1929. 10.1093/bioinformatics/btz795.

65. Edgar, R.C. (2004). MUSCLE: multiple sequence alignment with high accuracy and high throughput. Nucleic Acids Res 32, 1792–1797.

66. Jumper, J., Evans, R., Pritzel, A., Green, T., Figurnov, M., Ronneberger, O., Tunyasuvunakool, K., Bates, R., Zidek, A., Potapenko, A., et al. (2021). Highly accurate protein structure prediction with AlphaFold. Nature 596, 583–589. 10.1038/s41586-021-03819-2.

67. Soding, J. (2005). Protein homology detection by HMM-HMM comparison. Bioinformatics 21, 951–960. 10.1093/bioinformatics/bti125.

